# Assessing cellular metabolic dynamics with NAD(P)H fluorescence polarization imaging

**DOI:** 10.1101/2025.07.28.667273

**Authors:** Lu Ling, Jack C. Crowley, Matthew L. Tan, Jennie A.M.R Kunitake, Adrian A. Shimpi, Rebecca M. Williams, Lara A. Estroff, Claudia Fischbach, Warren R. Zipfel

## Abstract

Altered metabolism enables adaptive advantages for cancer, driving the need for improved methods for non-invasive long-term monitoring of cellular metabolism from organelle to population level. Here we present two-photon steady-state fluorescence polarization ratiometric microscopy (FPRM), a label-free imaging method that uses nicotinamide adenine dinucleotide (phosphate) (NAD(P)H) autofluorescence as a functional readout of cellular metabolism. The method is simple to implement and operates an order of magnitude faster than the NAD(P)H-fluorescence lifetime imaging microscopy (FLIM) imaging modality, reducing cytotoxic stress while providing long-term monitoring capacity. FPRM enables high-resolution dynamic tracking of NAD(P)H signals with subcellular details and we have established a set of instrument-independent ratiometric parameters that correlates NAD(P)H signals with metabolic status during pharmaceutical and environmental perturbations. We further integrated FPRM readouts with other parameters such as cell shape and migration on 2D and 3D collagen matrices, demonstrating the technique’s versatility across bioengineered platforms for cancer metabolism research.

## Main

Metabolic reprograming is a pan-cancer hallmark as cancer cells adapt their bioenergetic and biosynthetic programs to evade microenvironmental stresses and metastasize^1,2^. Our current understanding of how intrinsic changes in tumor cell metabolism affect cancer cell migratory and invasive capabilities, the first step of metastasis, remains limited, due in part to a lack of experimental approaches that allow for long-term monitoring of metabolic dynamics under relevant microenvironmental conditions. Despite their important contributions to cancer research, current state-of-the-art metabolomics techniques are often limited to endpoint measurements of cell populations^3^. For single cell details, fluorescence microscopy using genetically encoded or exogeneous fluorescent sensors of metabolites^4^ are commonly used to monitor metabolite dynamics. However, while exogenous sensors can be minimally invasive, they may perturb cellular functions either by acting as scavengers (“sponge effect”) or by interacting with other cellular components. For this reason, label-free imaging of nicotinamide adenine dinucleotide (phosphate) (NAD(P)H) autofluorescence has become an attractive method to assess cellular metabolic state^5^. The reduced forms of nicotinamide adenine dinucleotide (NAD(P)H) emit blue autofluorescence (460 nm peak), while the oxidized form is non-fluorescent^6^. Changes in cytosolic and mitochondrial NAD(P)H autofluorescence are taken to reflect the dynamics of glycolysis and oxidative metabolism, respectively^7^ since NADH does not diffuse passively through the mitochondrial membrane. Since NADH concentrations are ∼10 times higher than NADPH in most tissues^8^, researchers often refer to NADH as the primary fluorophore during imaging, and we will adapt the same approach here.

Early NADH imaging measurements were intensity based, at first using UV excitation^6^, and later using 700-750 nm two-photon excitation^9,10^, to sense changes in the redox balance and enzyme binding. However, intensity alone can lead to ambiguous results since it is non-ratiometric and sensitive to the excitation intensity, concentration, binding-state of the fluorophore^6^, and light scattering within 3D culture platforms. Fluorescence Lifetime Imaging Microcopy (FLIM) is independent of intensity measurements and thus a better imaging mode for NADH imaging. The fluorescence lifetime of NADH shifts between 0.2–0.4 ns in the unbound (or “free”) population to 1–2 ns for the protein bound population. Currently NADH-FLIM has been used to monitor cellular metabolism during stem cell differentiation, evolution of cancer cell heterogeneity, macrophage polarization and T cell activation^11–17^. While this method has great utility in single-cell metabolism research, it does have drawbacks due to instrumentation complexity and cost, and its long acquisition times (∼1–4 minutes/frame) limiting longitudinal studies^18^. The low two-photon action cross-section of NADH (∼0.08–0.25 GM)^19^ makes high throughput FLIM challenging, even with recent acquisition hardware improvements (see Supplementary Information section SI-1). More importantly, we have found that the long exposure times of what is essentially UV excitation within the focal plane perturbs cell metabolism, prohibiting longitudinal experiments (SI-3).

Similar to FLIM, time-resolved polarization anisotropy of NADH is independent of intensity and has been used by a number of groups^20–23^. Measuring the time-resolved rotational properties of free vs. bound NADH can provide more detailed information on the state of NADH compared to standard NADH-FLIM, but still requires long acquisition times. However, the rotational mobility is already encoded by the steady state polarized emissions, and time-resolved fluorescence methodologies are not necessary to assess NADH binding status. Two-photon imaging of steady state polarized emission is a simpler and faster approach that reduces phototoxicity and enables an expanded menu of potential applications. Here we present Fluorescence Polarization Ratiometric Microscopy (FPRM) as a robust method for analyzing free vs. bound NADH in live cells. FPRM uses a much shorter acquisition time per field of view (typically 1-5 seconds), significantly reducing phototoxic effects and allowing for higher temporal resolution NADH imaging. We describe the establishment of an imaging pipeline and define a set of system-independent ratiometric parameters to correlate NADH signals from subcellular organelle compartments to cellular metabolic status. We validated FPRM using metabolic inhibitors and physiological stresses such as hypoxia and demonstrate that it is a robust indicator of metabolic response to biologically relevant conditions. Lastly, we tracked tumor cell migration in 2D and 3D in vitro platforms, showing correlations between FPRM parameters and cell motility and shape. Our established imaging pipeline captures metabolic heterogeneity with subcellular resolution, enabling correlations of cell metabolism with time-dependent cellular responses to metabolic stressors as well as processes such as migration.

## Results

### Fluorescence Polarization Ratiometric Microscopy (FPRM) of NADH

Emission polarization from unbound or free NADH is more depolarized compared to protein-bound NADH. By measuring the steady state intensities of the perpendicular (S) and parallel (P) emissions, the extent of depolarization can be determined (Fig. 1a). Only minor modifications are required to enable emission polarization ratiometric imaging on a multiphoton laser scanning system (Fig. 1b).

**Figure 1.**
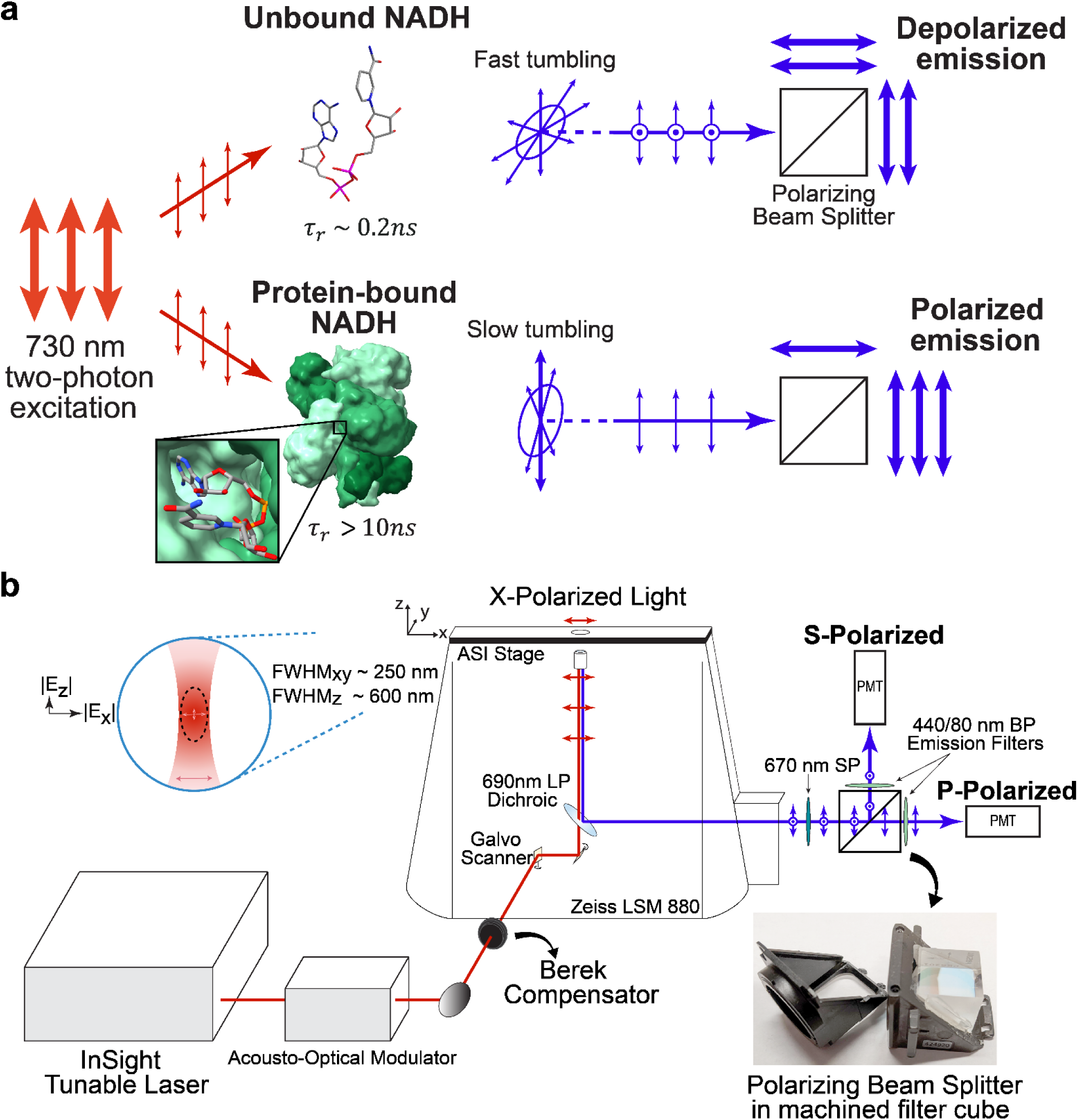
Steady-state Fluorescence Polarization Ratiometric Microscopy (FPRM). **(a)** The fluorescence emission from protein-bound NAD(P)H is significantly more polarized than emission from unbound NAD(P)H. (top) NADH in aqueous environments has a rotational correlation time on the order of 0.2 ns, while (bottom) the rotational correlation time of globular proteins that bind NADH can be orders of magnitude slower. **(b)** P-polarization refers to emissions polarized parallel to the excitation polarization, while S-polarization refers to emissions perpendicular to the excitation polarization. Detecting differences in steady-state polarization is carried out using a polarizing beam splitter placed in a modified filter cube which directs the S and P polarized emissions to two GaAsP PMTs in the non-descanned detection path. Identical emission filters (440/80 nm bandpass “BP”) were used to collect NAD(P)H signals. A 670-short pass (“SP”) filter was placed before the beam splitting cube to further suppress the 730 nm excitation. Linear polarization of the 730 nm excitation at the objective back aperture was obtained using a Berek Compensator to compensate for polarization distortions caused by the coupling optics and galvo mirrors. Blue inset: a high-numerical aperture (NA) objective lens enables focal point excitation with several hundred nm axial and lateral resolution. Typical point spread function full width at half-maximum “FWHM” values for imaging in this work are listed. One consequence of high-NA optics is the appearance of a considerable z-axis component to the light polarization, as we discuss in Supplemental Information section SI-2.3.

Both fluorescence lifetime and steady state emission polarization are sensitive to the ratio of unbound NADH to bound NADH, and both eliminate errors that can occur with intensity-only measurements. To demonstrate that FPRM detects free vs. bound NADH with comparable sensitivity to NADH-FLIM, we measured the NADH polarization ratio, fluorescence intensity, and fluorescence lifetime measurements of a solution of NADH titrated with varying concentrations of the NADH-binding protein glutamate dehydrogenase (GDH). Figure 2a and 2b show the change in the emission channel intensity and P/S polarization ratio with increasing concentrations of GDH-NADH binding sites. The increase in brightness upon binding depends on the protein to which it binds (SI-1). The observed binding kinetics can be modeled (Fig. 2b, SI-2.2) and returns a K_D_ value for NADH binding of 26 µM, within the range of reported KD’s for NADH binding which vary between 3.2 to > 100 μM^24,25^. Figure 2b also shows the change in lifetime in the same samples, which similarly correlates with binding.

**Figure 2.**
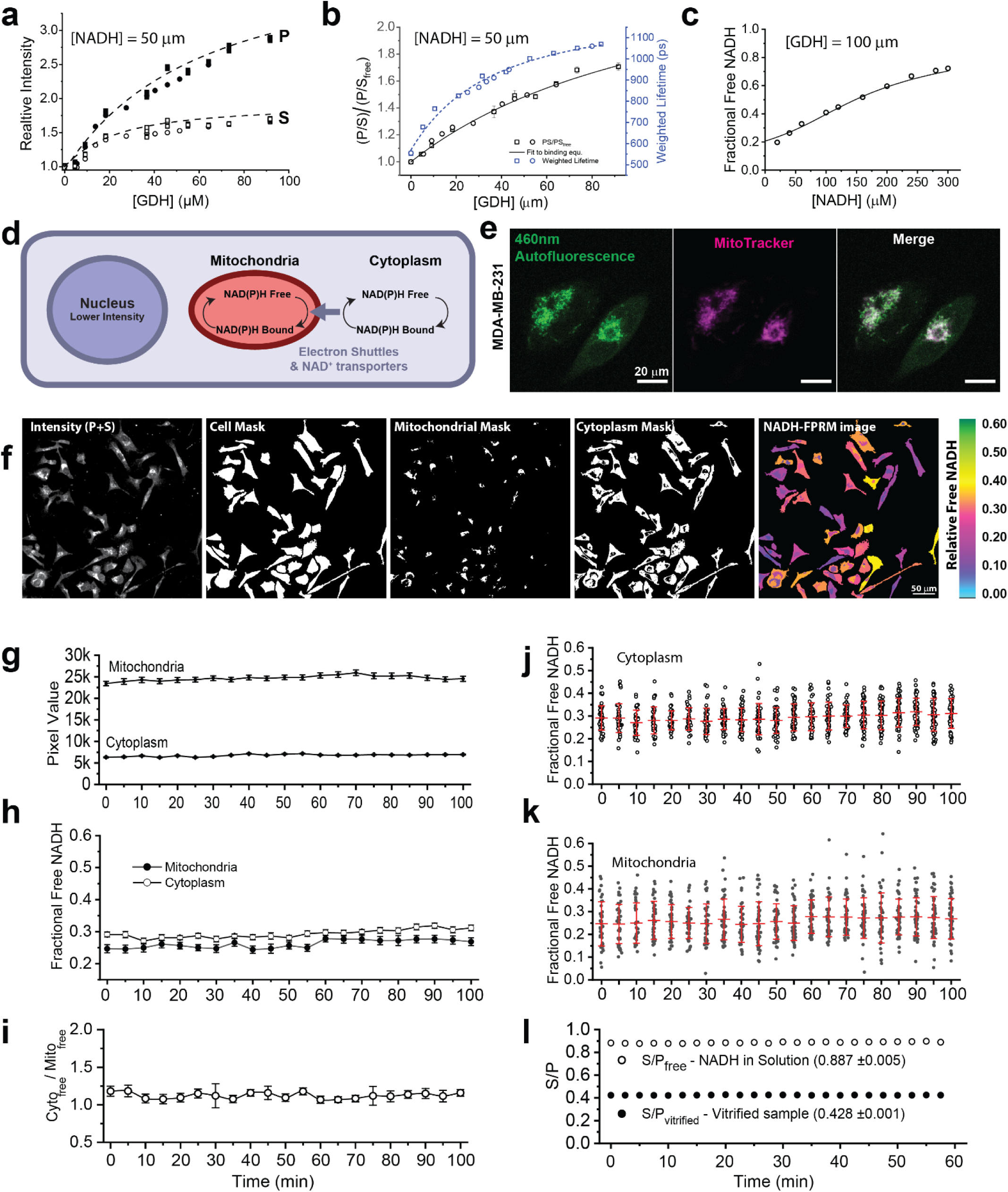
FPRM imaging of NAD(P)H. **(a)** Relative S and P intensities from an *in vitro* titration of NADH (50 μM) with increasing concentrations of GDH. **(b)** P/S ratio and weighted fluorescence lifetimes (2-exp fits) as a function of [GDH]. Different symbols in (a) and (b) are results from two separate experiments. Dashed lines in (a) and (b) are polynomial fits to show the trend; solid line in (b) is results from the binding model described in the SI-2.2. **(c)** Fractional free NADH calculated based on the S/P ratio and equation 1 with increasing concentrations of NADH in the presence of 100 μM GDH. Solid line is from a binding model described in SI-2.2. **(d)** Schematic showing subcellular distribution of NAD(P)H in cells. NAD(P)H is partitioned into three cellular “compartments,” from which exchange occurs through various shuttle systems. **(e)** MitoTracker Red fluorescence and endogenous fluorescence signal used in FPRM shows spatial colocalization in MDA-MB-231 cells. **(f)** FPRM analysis pipeline: simultaneously acquired images of P- and S-polarized emissions are collected. A summed total intensity (P+S) image is used to generate cell, mitochondrial, and cytoplasmic masks by thresholding. The masks are then applied to the S and P images to obtain S and P pixel values from ROIs from each compartment. Relative free NAD(P)H levels are calculated as [(S/P) – (S/P) _vitrified_] / [(S/P) _free_ – (S/P) _vitrified_] and compartmentalized FPRM color-coded images can be created. **(g-k)** 100-minute time course measurements of MDA-MB-231 cells. **(g)** Mean pixel intensity values from the mitochondrial and cytoplasmic compartments from 46 cells (at each time point). **(h)** Relative fractional levels of free NAD(P)H in the mitochondrial and cytoplasmic compartments, and **(i)** Cyto_free_/Mito_free_ NAD(P)H_free_ ratios. Values in **(g)** – **(i)** are means ± SEM acquired at 5-minute intervals. Images were acquired using 730 nm delivered through a 63x/1.4 NA objective lens (∼6 mW, ∼120 fs pulse width, four-frame average at 1.2 s/frame). **(j, k)** The large deviations between cells at each time point reflects actual metabolic differences and are not measurement noise. This metabolic state heterogeneity can also be seen in the color-coded image in **(f)**. For comparison **(l)** shows S/P values from a NADH in solution and from vitrified sample (blue fluorescent slide) which show minimal deviations. S/P values shown in **(l)** define the limits used to normalize the S/P cell data for images acquired using the 63x/1.4 Zeiss Plan-Apochromat objective lens.

When the NADH concentration exceeds the concentration of NADH binding sites, the S polarized emission increases as the fraction of unbound (free) NADH increases. Using the measured S/P ratio the unbound fraction of NADH can be estimated by normalization between measurable S/P limits:

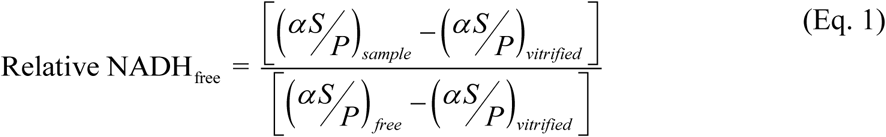

α is a prefactor (α·S/P) which varies between objective lenses, photomultiplier tubes (PMTs) and the light path of different microscopes. The prefactor α is canceled in the ratio equation above, eliminating imaging system instrument dependence. Based on equation 1 the fraction of free NADH can be calculated, as illustrated is in Fig. 2c which shows the increase in free NADH as a function of varying [NADH] (20 to 300 μM) at a constant (100 μM) GDH-NADH binding site concentration.

We note that our fluorescence polarization normalization (Eq. 1) differs from the steady state fluorescence anisotropy normalization commonly used in fluorescence spectroscopy^26^ or in implementations of time-resolved lifetime anisotropy measurements^21^, and our approach is further justified in the supplementary information section (SI-2).

NAD(P)^+^/NAD(P)H pools are present in different cellular compartments including the nucleus, mitochondria, and cytoplasm and involved in metabolism, redox balance, and gene expression^27^ (Fig. 2d). Based on measurements and estimated ranges for concentrations of NADH and NADH binding sites from the literature (see Appendix SI-Table 2), the NADH concentration typically exceeds that of binding sites, and we find that FPRM and equation 1 provides a simple means to monitor changes in the amount of unbound NADH in cellular compartments. Detecting changes in the relative amounts of free NADH in the cytoplasm and mitochondria is invaluable, since each has a distinct, but direct relationship to cellular metabolism.

We examined the use of FPRM in MDA-MB-231 cells, an extensively-characterized, highly aggressive breast cancer cell line associated with altered metabolism^28^. We use segmentation to separate NADH dynamics in the mitochondrial compartments from the cytoplasmic region, providing two compartment-based metrics that correlate with changes in the fraction of unbound NADH in each compartment, which in turn is an indicator of the metabolic state of the cell. To justify using intensity-based image segmentation for label-free separation of mitochondrial signals from the cytoplasm we used MitoTracker-Red to validate that the higher intensity regions colocalized with mitochondria (Fig 2e). As expected, these regions within MDA-MB-231 cells co-localized with mitochondria, as others have shown^5^. Because nuclear and cytoplasmic compartments have a faster means of metabolite exchange between them compared to the slower mitochondrial shuttles^29–31^, we include the nuclear NADH signal as part of the cytoplasmic pool.

The pipeline for image analysis is illustrated in Figure 2f. Polarized emission images from S and P channels are simultaneously collected during live imaging and the S and P pixel intensities are summed to provide a total intensity image, which is used to create cytoplasmic and mitochondrial region masks by thresholding. The cytoplasm mask is generated by subtracting the mitochondria mask from the whole cell mask.

The average intensity of S and P per cell within each region-of-interest (Fig. 2g) is used to calculate S/P values, which are then normalized between free NADH in solution (PBS) and a vitrified sample (blue plastic slide) to calculate the fractional unbound NADH based on equation 1. Our fractional free NADH estimate is expected to be slightly underestimated since no protein immobilizes NADH to the same extent as the vitrified sample we use for (S/P)_vitrified_. We chose to use a normalization standard that is easy and robust, rather than using a specific NADH-protein mixture, since the intracellular NADH binding conditions are complicated and vary between cell type and cellular compartments. Using a simple, standardized normalization method sample is critical for making repeatable measurements over time and over different instruments (see SI-2.3).

In MDA-MB-231 cells under standard culture conditions, we find that average fractional free NADH values from mitochondria and cytoplasm compartments are typically in the 0.2 – 0.35 and 0.25 - 0.45 range, respectively (Fig. 2h). Using FPRM we can acquire long time courses (e.g., Fig. 2g–i) without any obvious changes in FPRM parameters, indicating minimal metabolic perturbation at the populational level. Interestingly, even under steady state culture conditions, free NADH values of individual cells fluctuate over time for both cytoplasmic and mitochondrial regions (Fig. 2j-k). These fluctuations are due to instantaneous physiological/metabolic differences between single cells, rather than measurement artifacts (e.g. noise) from the imaging system, as S/P_free_ and S/P_vitrified_ remained stable (Fig. 2l). Cancer cell metabolism is inherently heterogeneous and dynamic, and our data suggest that FPRM captures these changes at the single-cell level with subcellular resolution without perturbing population-level behaviors during long-term imaging.

Overall, we have found that the FPRM parameters that estimate the fractional free NADH (Mito_free_, Cyto_free_) and the ratio Cyto_free_/Mito_free_ are robust indicators of cellular metabolism that are faster and require simpler instrumentation, compared to NADH-FLIM. The metrics are insensitive to laser intensity differences, microscope and objective lenses used, as well as relatively insensitive to threshold levels used to separate compartments (SI-2.3).

To validate that FPRM imaging is less metabolically stressful than FLIM due to a lower irradiation dose, we examined oxidative stress as a function of continuous irradiation times ranging from the typical acquisition condition used in FPRM to the longer acquisition times used for NADH-FLIM (Fig. SI-3). To determine oxidative stress, we incubated cells with biotinylated glutathione ethyl ester (BioGEE), which allows detection of protein glutathionylation by endogenous glutathione reductase (GR) in the presence of reactive oxygen species^32^, followed by quantification using a fluorophore labeled streptavidin (Fig. SI-3a-c). Using this strategy, we carried out irradiations at 700 nm of cells for 60, 90 and 240 scans to mimic typical FLIM acquisition doses, and 4 scans to represent 4-line average frame in our typical FPRM acquisition mode. We observed that 60-240 continuous scans resulted in a measurable increase in protein labeling in the mitochondria, compared to non-irradiated cells, indicating an increased generation of reactive oxygen species (Fig. SI-3d). The shorter 4 second scan average irradiation used to emulate NADH FPRM measurements did not produce significant changes in glutathionylation levels, suggesting that NADH polarization imaging at a few seconds per frame induces significantly less oxidative stress than NADH-FLIM, which requires longer integration times per frame. We next assessed whether prolonged scanning with 730 nm pulsed excitation produced changes in respiration by monitoring oxygen uptake after imaging (Fig. SI-3f-h). We saw increases in the respiration rate when measured after 200 seconds of scanning at 730 nm (10 mW delivered through a 1.0 NA objective). These measurements indicate that prolonged continuous scanning at in the 700 nm range at powers needed for two-photon NADH excitation can perturb cellular metabolism by generating oxidative stress. The lower excitation dosage required for FPRM greatly reduces the extent of metabolic perturbation compared to typical NADH-FLIM measurements.

### Artifacts from autofluorescent non-NAD(P)H containing compartments

To expand the use of FPRM to tumor cells with varying levels of malignant potential, we examined isogenic cell lines from the MCF10A series: MCF10A (non-transformed mammary epithelial cells) and its aggressive counterpart MCF10CA1a (metastatic *in vivo*)^33^ (Fig 3a). While the brighter autofluorescence regions in MCF10A cells co-localized with MitoTracker-labeled mitochondria, MCF10CA1a cells additionally contained autofluorescent granules that did not co-localize with mitochondria (Fig. 3a). Emission spectra of the granules showed a red-shifted emission compared to NADH in solution or in mitochondria (Fig 3b). FLIM of MCF10CA1a cells additionally detected longer amplitude-weighted lifetimes in these granules (>1.8 ns) compared to mitochondrial regions (Fig 3c).

**Figure 3.**
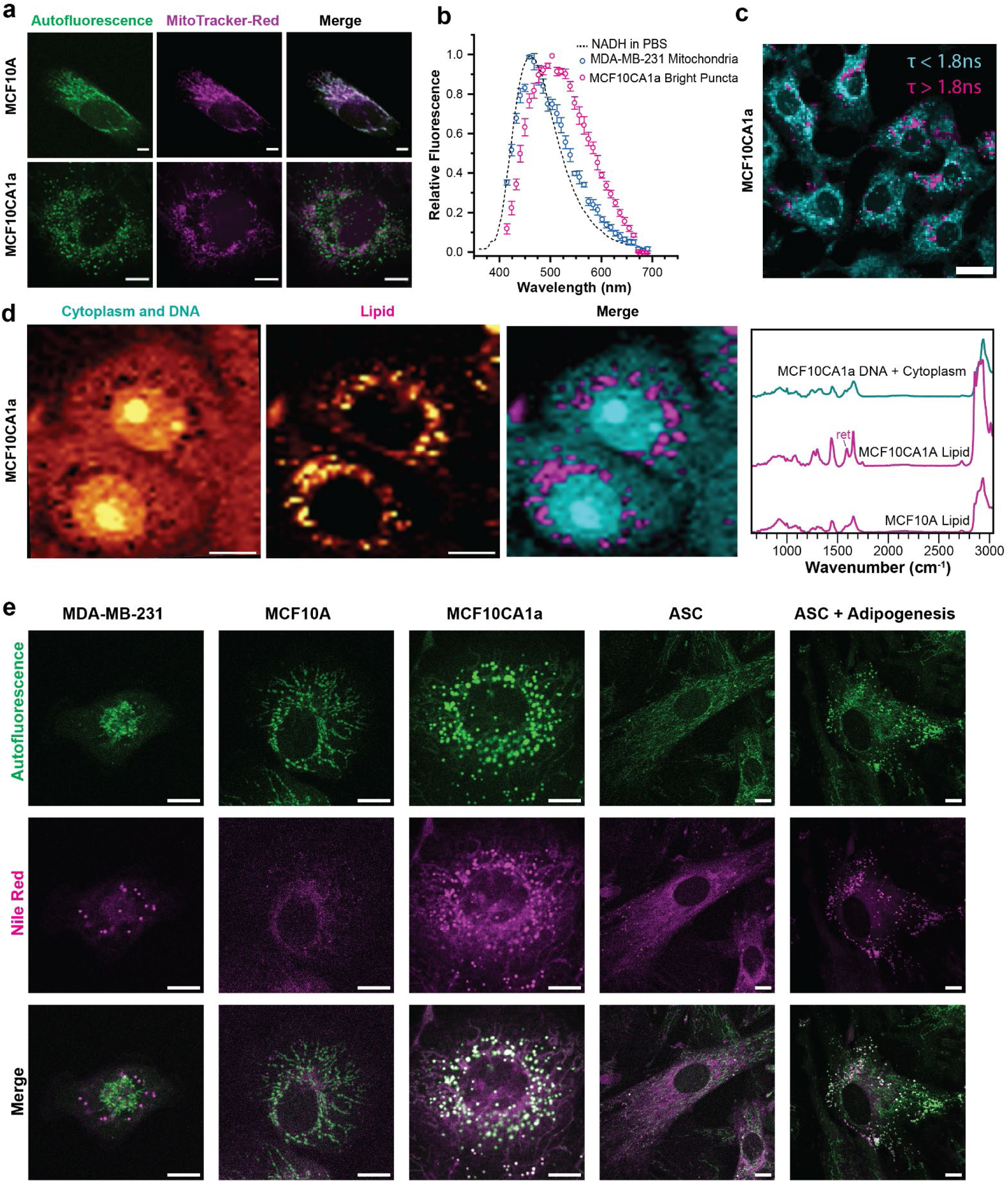
460 nm Autofluorescent structures need to be validated to perform analytical NAD(P)H fluorescence microscopy. **(a)** Representative overlaid images of 460 nm autofluorescence intensity (green) and MitoTracker Red fluorescence (magenta), demonstrating that brighter fluorescence signals of NAD(P)H colocalize with mitochondrial structures in MCF10A cells (as in MDA-MB-231 cells in Fig. 2), while not all bright puncta colocalize in MCF10CA1a cells. **(b)** 730 nm excited autofluorescence emission spectra from NADH in solution, mitochondrial regions in MDA-MB-231 cells, and bright autofluorescence puncta in MCF10CA1a cells. NADH spectrum measured in a single trace; MDA-MB-231 and MCF10CA1a circles are means ± s.d. (n=3). **(c)** Intensity image from FLIM acquisition of MCF10CA1a cells, where a 2-color encoding indicates pixel amplitude-weighted lifetimes that were below (cyan) or above (magenta) a 1.8 ns threshold. Bright spots in some cells are above the 1.8 ns lifetime threshold. **(d)** Representative MCF10CA1a confocal Raman microscopy false color images (left), overlaying DNA/cytoplasm (blue) and lipid (magenta) signatures, and corresponding Raman spectral averages (right), with lipid-rich spectral average for MCF10A shown for comparison. Peak at 1592 cm^-1^ labeled “ret” corresponds to a retinol-like signature. **(e)** Representative overlaid images of 460 nm autofluorescence intensity (green), and Nile red stain of lipid droplets (magenta) in various cell lines.

In an effort to determine the identity of the autofluorescent granules and because we have previously shown that MCF10CA1a contain increased levels of cytoplasmic lipid compared to their less aggressive counterparts^34^, we analyzed MCF10CA1a by Raman microscopy to assess the chemical composition of subcellular structures^35^. Interestingly, this approach confirmed the presence of lipid-rich vesicular structures suggesting that the autofluorescent granules contain lipid (Fig. 3d). False color Raman peak area images enable the visualization of the spatial distribution of biological components based on integration of peaks characteristic to the distinctive chemistries of certain classes of biological molecules^36,37^. False color images corresponding to DNA/cytoplasm signatures (by integrating from 2928-2988 cm^-1^) and lipid signatures (by integration of the characteristic peak at 2850 cm^-1^) are shown individually and merged (DNA/cytoplasm-rich in blue and lipid-rich in magenta) for representative cells of MCF10CA1a. Corresponding average spectra from the highest intensity pixels are shown (DNA/cytoplasm-rich average (blue trace) and lipid-rich average (magenta traces)), and the lipid-rich average spectrum for MCF10A is shown for comparison. The lipid-rich averages contain peaks characteristic of lipids (including 2850 cm^-1^, and 1300 cm^-1^ among others^36^). Notably, the granules in MCF10CA1a cells also contain a distinct, sharp peak at 1592 cm^-1^ which was not observed in lipids in MCF10A (Fig. 3d and SI-4). This may be linked to a retinol- or retinyl ester-like signature that overlaps with the NADH fluorescence spectrum^38^, which could explain the autofluorescence observed in the MCF10CA1a cells during FPRM. Using Nile Red staining for neutral lipids, we confirmed that the autofluorescent granules in MCF10CA1a cells co-localized with neutral lipid droplets (Fig. 3e). Interestingly, Nile Red staining suggested that MDA-MB-231 cells also contain lipid droplets, but these structures were not autofluorescent in the 400-500 nm range. To confirm that cytoplasmic lipid can lead to autofluorescence in the NADH emission range, adipose stem cells (ASC) were induced to undergo adipogenesis. Indeed, post-differentiation ASCs accumulated lipid droplets whose autofluorescence was consistent with NADH emission (Fig. 3e). While the generation of lipid bodies may be associated with altered metabolism, it is important to correctly attribute autofluorescence to NADH sources when probing cellular metabolic status, and investigators should always check for lipid-based artifacts and mitigate any contamination by segmentation when using FPRM (or FLIM). For all subsequent FPRM experiments in this paper, we utilized only MDA-MB-231 cells in which 460 nm autofluorescent lipid granules were absent.

### FPRM is a robust metric for cellular metabolic status

To validate that measured FPRM parameters correlate with dynamic changes in NADH state, and determine how those changes are related to metabolism, we subjected MDA-MB-231 cells to various metabolic perturbations, such as potassium cyanide (KCN) respiration inhibition (Fig. 4a), carbonyl cyanide 4-(trifluoromethoxy)phenylhydrazone (FCCP) uncoupling of oxidative phosphorylation (Fig. 4b), 2-deoxy-D-glucose (2DG) glycolysis inhibition (Fig. 4c), and AZD7545 pyruvate dehydrogenase complex (PDC) inhibition, which stimulates oxidative phosphorylation (Fig. 4d). Using FPRM, we observed cells in an untreated state for 15 minutes before initiating treatment, and continuing acquisition for 30-50 minutes. Cells were imaged at 5-minute intervals, and the changes in the S and P cytoplasmic and mitochondrial compartment intensities were monitored over time. Analysis was carried out on an individual cell basis and population mean ± SEM at each time point were plotted (Fig 4a-d). The heterogeneity in FPRM measurements between cells is clearly represented by large standard deviations. Heat maps of individual cell responses to KCN, FCCP, 2DG and AZD7574 shown in SI-5 further illustrate cell-to-cell heterogeneity, where each cell responds uniquely to the same treatment.

**Figure 4.**
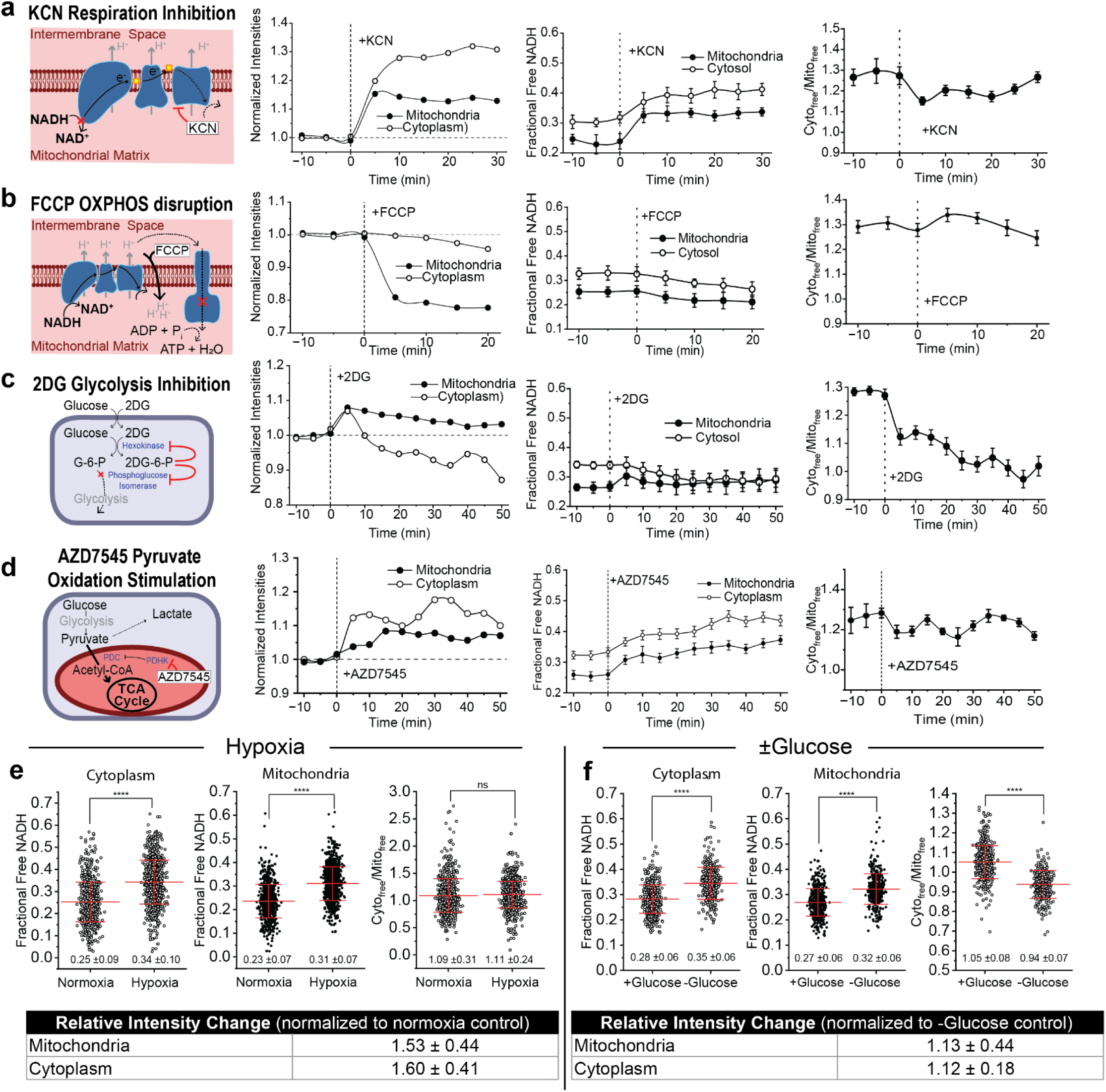
NAD(P)H polarization ratio dynamics correlate with pharmaceutically induced metabolic alterations and environmental stresses in MDA-MB-231 cells. **(a)-(d)** FPRM response to the common metabolic inhibitors KCN (5 mM), FCCP (5 μM), 2DG (50 mM) and ADZ7545 (5 μM). Each row shows an illustration of the mode of action followed by plots of FPRM time course measurements of MDA-MB-231 cells after adding the inhibitor. The P and S normalized mean fluorescence intensities, fractional free NADH in the cytoplasmic and mitochondrial compartments, and the ratio of the compartment values (Cyto_free_/Mito_free_) are plotted. All data was taken at using 730 nm excitation with ∼6 mW at the focus delivered through a Zeiss Plan-Apochromat 63x/1.40 objective. The mean fractional free NADH values for the field of cells were calculated using equation 1 with values of 0.89 for S/P_free_ and 0.43 for S/P_vitrified_. Error bars are the standard error of the mean (SEM). Average number of cells per time point (cells migrate into and out of the field): KCN data is from ∼60 cells, FCCP data from ∼90 cells, 2DG from ∼80 cells, and AZD7545 from ∼120 cells. Abbreviations used in illustrations: ADP=Adenosine diphosphate, ATP=Adenosine triphosphate, G-6-P=Glucose-6-phosphate, 2DG-6-P=2-Deoxy-D-glucose-6-phosphate, PDHK=Pyruvate dehydrogenase kinase, PDC=Pyruvate dehydrogenase complex, TCA=Tricarboxylic acid cycle. **(e)** Effect of hypoxia on FPRM parameters. Scatter plots: FPRM measurements from MDA-MB-231 cells following 3 days of incubation in normoxic (20% O_2_, data from 475 cells) vs. hypoxic (1% O_2_, 658 cells) environments. Circles represent measurements in individual cells. Red lines are the mean and SEM (whisker lines). Table: relative summed intensity (S+P) changes in cytoplasmic and mitochondrial compartments. **(f)** Effect of glucose deprivation on FPRM parameters in MDA-MB-231 cells. Cells were moved from the standard +glucose media to glucose free media and imaged after 30 minutes. Scatter plots: free NAD(P)H in cytoplasmic and mitochondrial regions and the relative summed intensity changes from the two compartments. Circles represent measurements in individual cells (531 cells in +glucose media and 420 cells after 30 minutes in glucose free media). Red center lines are the mean and whisker lines are the standard deviation.

KCN blocks electron transport, stopping NADH oxidation in mitochondria which leads to an accumulation of free NADH^11^. As expected, FPRM time course measurements (Fig. 4a) show an increase in free NADH from cytochrome C inhibition. Unbound NADH fraction increases in both mitochondrial and cytoplasmic compartments, with mitochondrial free NADH increasing to a larger extent (40% vs. 35%), resulting in a slight decrease in the Cyto_free_/Mito_free_ ratio.

The uncoupler FCCP escorts protons across bilipid membranes, eliminating the inner mitochondrial membrane proton gradient that drives ATP phosphorylation^39^ (Fig. 4b). Upon treatment, FCCP induced a rapid drop in mitochondria intensity and a slight decrease in cytosol, similar to other reports demonstrating that FCCP enables continuous oxidation of mitochondrial NADH^40^. In contrast, the average fractional free NADH remained relatively stable with a slight decrease in both mitochondria and cytosol. Cyto_free_/Mito_free_ exhibited an initial increase post treatment and a gradual return to new equilibrium. This is consistent with other work showing FCCP treatment led to a robust increase in NAD^+^/NADH ratio, even in the cytosol, suggesting maximal respiration consumes both mitochondrial and cytoplasmic NADH pools^40^.

2DG inhibits the production of glucose-6-phosphate from glucose via hexokinase and phosphoglucoisomerase interference and therefore inhibits glucose catabolism^41^ (Fig. 4c). In comparison to controls, a brief spike and then a continuous decrease in fluorescence intensity was observed in both mitochondrial and cytoplasmic regions, presumably due to reduced flux into both glycolysis and the TCA cycle, which supports NAD^+^/NADH homeostasis. The cell averaged fractional free NADH increased in cytosol while decreased in mitochondria, resulting in a decrease in Cyto_free_/Mito_free_ (Fig. 4b). Since MDA-MB-231 cells are aggressive cancer cells that rely on aerobic glycolysis for metabolism, inhibiting glycolysis likely leads to depletion of free NADH in cytoplasm, and increased fraction free NADH in cytoplasm as cells switch to oxidative phosphorylation (OXPHOS)^42^, potentially prioritizing glutamine catabolism, to compensate.

PDH regulates the first step of pyruvate conversion to acetyl-CoA (Fig. 4d) and has been linked to changes in the NAD^+^/NADH ratio within cells which competes with ATP production^43^. AZD7545 inhibits PDH complexes and promotes the shifting of metabolic preferences to OXPHOS in various cell lines, however the expected changes in NAD(P)H are not clear^43^. Upon AZD7545 treatment, the mitochondrial NADH intensity increased more drastically compared to the cytoplasmic signal (Fig. 4c), suggesting that there was a rapid increase in the accumulation of both free and protein-bound NADH fractions, consistent with the observation of decreased NAD^+^/NADH in cells as previously reported^43^. Cyto_free_/Mito_free_ decreased toward unity in response to both 2DG and AZD7545 treatments, sensing considerably different metabolic stresses. In 2DG, a loss of glycolysis and TCA functionality appears to deplete the NADH pool in both compartments, leaving a low, mostly-bound NADH population with low cytoplasmic vs. mitochondrial polarization contrast. In contrast, AZD7545 shows lost cytoplasmic vs. mitochondrial contrast as uncompensated NAD^+^ reduction by glycolysis and TCA activity leads to over-saturated free NADH levels in both compartments.

The deviations observed in these time course experiments are due to cell-to-cell variations in metabolic state in response to the inhibitor. This can be seen in the heatmaps of FPRM parameters from individual cells plotted vs time after addition of the drug treatments, in which some cells show dramatic changes in Cyto_free_/Mito_free_, while others show relatively little change (SI-5, figures a-d). These data highlight intrinsic metabolic heterogeneity within a population where cells may adapt uniquely in response to metabolic stress.

To broaden our comprehension of FPRM-measured NADH dynamics in response to more complex metabolic alterations, we exposed MDA-MB-231 cells to different environmental stressors, including hypoxia (Fig. 4e). After 3 days of restricted oxygen levels (1%), cells have more drastically increased cytosolic fraction free NADH and mitochondrial fraction free NADH, leading to an unchanged Cyto_free_/Mito_free_ compared to normoxia (20%). Intensities in both the cytoplasmic and mitochondrial compartments had a ∼50% increase compared to normoxia, suggesting a shift in NAD^+^/ NADH ratio due to a slower oxidation rate in the electron transport chain. Excessive buildup of NADH is a classic signature of hypoxia and cancer cells can adapt to serine catabolism to maintain NADH levels for sustained cell proliferation^44^

As starvation of nutrients limits metabolic fuels entering various metabolic pathways, we examined the metabolic responses of MDA-MB-231 cells following removal of glucose using FPRM. Loss of glucose limited metabolic fuels entering glycolysis and TCA cycles, and cells in glucose-excluded media displayed a significantly higher fractional free NADH in both mitochondria and cytosol regions compared to cells in glucose containing media (Fig. 4f). Similar to 2DG treatment, Cyto_free_/Mito_free_ in glucose-excluded conditions were significantly lower than glucose conditions.

Overall, FPRM parameters such as the normalized compartmental intensities, free cytoplasmic and mitochondrial NADH, and the Cyto_free_/Mito_free_ ratio provided powerful and robust measurements of cellular metabolism dynamics. The comparative changes in these readouts offer strong indicators of metabolic shifts as cells respond and adapt to different stressors.

### FPRM of cell motility in 2D

Invasion of cancer cells into the surrounding host tissue is the first step of metastasis, the leading cause of death for patients with advanced disease. This migratory behavior is an energy-consuming process, and a better understanding of the underlying metabolic dynamics could provide valuable insights for cancer research. After validating the feasibility and reproducibility of FPRM to understand NAD(P)H dynamics as an indicator of cell metabolism, we next used FPRM to determine how NADH dynamics correlate with individual cell motility. First, we performed these measurements in conventional 2D cell culture, and representative examples of a slow- and a fast-moving MDA-MB-231 cell are shown in Figure 5a-b.

**Figure 5.**
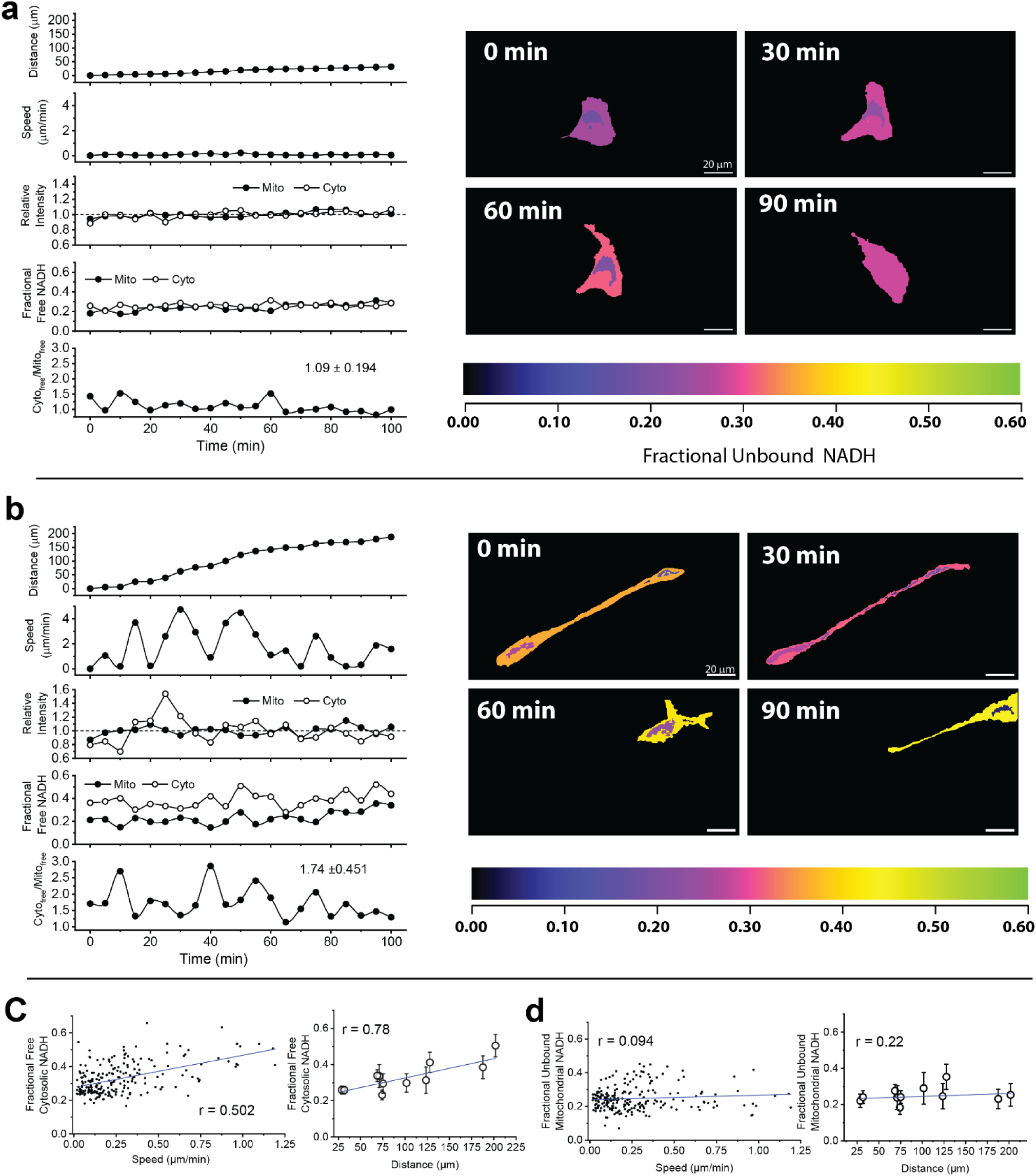
FPRM measurements correlate with migratory potential in 2D culture. Representative FPRM measurements for **(a)** a slowly moving cell, and **(b)** a rapidly migrating cell, over the course of 100 minutes. Time traces depict the total displacement of the cell’s centroid relative to time=0 min (“Distance”), the displacement of the cell centroid from frame to frame divided by the 5-minute acquisition interval (“Speed”), the normalized intensity change in the two compartments (“Relative Intensity”), the fractional unbound NADH levels for the mitochondrial and cytoplasmic compartments, and the ratio of those values (“Cyto_free_/Mito_free_”). **(c, left graph)** Cell migration speed (μm/min between time points) vs. fractional free cytoplasmic NADH measured during a 100-minute time course FPRM imaging of MDA-MB-231 cells. **(c, right graph)** Total distance traveled for 11 tracked cells vs cytoplasm unbound NADH (mean ±standard deviation). **(d, left graph)** Migration speed vs fractional free mitochondrial NADH measured during the time course; (**d, right graph) t**otal distance traveled vs mitochondrial free NADH (mean ±standard deviation).

While the slow cell remained relatively immobile, showing minimal changes in speed and migration distance, the fast cell readily altered its shape and migration speed as it traversed a steadily extending migration path (Fig. 5b). The relative intensities in both the mitochondrial and cytosolic compartments remained consistent in the slow cell, whereas the cytosolic intensity in the fast cell fluctuated dynamically, with minimal changes in the mitochondria (Fig. 5a). Interestingly, the fast-migrating cell exhibited a similar fractional free mitochondrial NADH value but a markedly increased fractional free cytosolic NADH (Fig. 5b). Cyto_free_/Mito_free_ in the fast cell also changed dynamically during migration, whereas the ratio remained largely stable in the slow cell. Time course videos of other representative stationary and migrating cells are presented in SI-6.

In a selection of unstressed cells representing a spectrum of migration potentials, we found that aggregate migration distance correlated linearly with fractional free cytoplasmic NADH instead of free mitochondrial NADH (Fig. 5c-d). As MDA-MB-231 cells are highly glycolytic^45^, cytoplasmic energy generation may be required to fuel migratory behavior, reflected in their dependence on free cytoplasmic NADH. The tracking capability of FPRM provided direct evidence linking migration to free NADH dynamics with compartment-specific metabolic preferences.

### Comparison of cell migration in 2D vs 3D engineered platforms using FPRM

While 2D culture experiments offer invaluable insights into cell behavior, they do not capture many of the biophysical aspects of the 3D microenvironment that cells navigate during invasion, such as extracellular matrix (ECM)-mediated changes in tissue architecture that lead to cell confinement^34^ and consequential changes in cell phenotype^46^. Indeed, invading breast cancer cells encounter an ECM that is particularly rich in collagen I^47,48^. Cancer cells can adhere to collagen via integrins and engage their actomyosin network to generate contractile forces that drive various cellular behaviors such as polarity, proliferation, migration, and malignant transformation^49^. ROCK activation is associated with cytoskeletal stress fiber formation leading to changes in cell shape associated with various modes of migration^50^. How ROCK signaling affects metabolism and if ROCK-dependent changes of metabolism differ between 2D vs. 3D culture remains unclear. To explore these relationships, we embedded MDA-MB-231 cells into 3D collagen hydrogels with defined stiffness and microarchitecture^46,51^. Additionally, we seeded cells on top of collagen-coated 2D glass substrates and added the ROCK inhibitor Y-27632^52^ to both culture contexts (Fig. 6a). Since FPRM image analysis uses segmentation of cell shape and time course tracking, we can simultaneously quantify NAD(P)H dynamics and other relevant parameters of cell state such as morphological changes and migration speed that are associated with ROCK inhibition.

**Figure 6.**
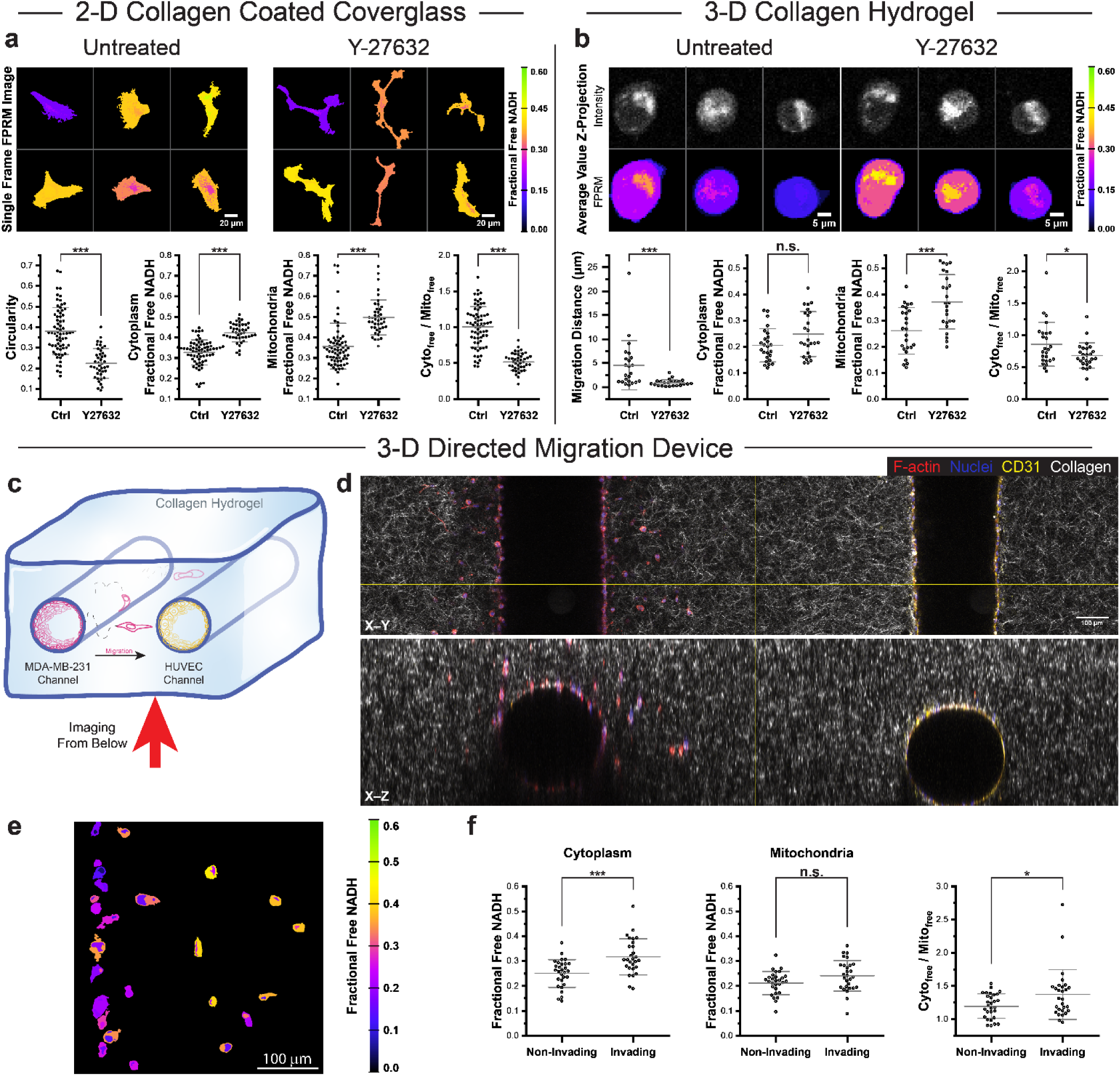
Application of FPRM in 2D and 3D engineered platforms. Comparison of MDA-MB-231 cell morphology and unbound NADH changes on (a) 2D collagen-coated coverglass and (b) within 3D collagen scaffolds, before and after inhibition of rho-associated protein kinase (ROCK) inhibition by Y-27632 treatment. **(a)** (Top) “Untreated:” color-coded FPRM images of representative cells on 2D collagen-coated coverglass, from time course FPRM after 30 minutes of steady-state imaging. “Y-27632:” color-coded FPRM images of the same cells after 60 minutes of Y-27632 treatment. (Bottom) Summarized results (circles represent measurements from individual cells) showing the shape descriptor of contour circularity (“Circularity”), fractional cytoplasmic and mitochondrial free NADH levels, and Cyto_free_/Mito_free_ ratios, between control (“Ctrl”) and Y-27632 treatment (30 μM). “Ctrl” shows the 68 measurements after the first 30 min of imaging before adding Y-27632. “Y-27632” shows the measurements of timepoints from 42 cells 30 min after adding Y-27632 at the 60-minute point. **(b)** (Top) “Untreated:” 5-slice average value Z-projection images of grayscale intensity (top) and color-coded FPRM (bottom) image stacks of 3 representative cells embedded in 3D collagen hydrogels, from time course FPRM after 30 minutes steady-state imaging. “Y-27632:” 5-slice average value Z-Projection images of grayscale intensity (top) and color-coded FPRM (bottom) image stacks of the same 3 cells after 150 minutes treatment with Y-27632. (below) Summarized results showing the 3D migration distance, fractional cytoplasmic and mitochondrial free NADH levels, and Cyto_free_/Mito_free_ ratios from a subset of cells before and after Y-27632 treatment. “Ctrl” shows FPRM measurements from timepoints after 30-45 min of imaging without the drug, while “Y-27632” shows the average measurements of timepoints 135-150 min after addition of Y-27632 treatment. **(c)** Schematic of the salient features of the engineered stroma directional migration device. Two parallel needle-size cylindrical lumens are formed in collagen hydrogels. One lumen was seeded with MDA-MB-231 cells, and the other was seeded with human umbilical vein endothelial cells (HUVEC) that form a vascular structure. Chemoattractants emitted by the HUVEC cells stimulate directed migration of tumor cells, which can be imaged by FPRM. **(d)** Immunofluorescence images of both endothelial and cancer cell lumens in 3D collagen scaffolds are shown in cross sectional and orthogonal views. Filamentous Actin (“F-actin”) staining with phalloidin is shown in red, nuclear staining with DAPI in blue, CD31 immunofluorescence in yellow, and confocal reflectance imaging of collagen is gray. **(e)** Color-coded FPRM image showing a subset of cells that have migrated towards the HUVEC channel. **(f)** Summarized FPRM measurements showing fractional free NADH in the cytoplasmic and mitochondrial regions, and Cyto_free_/Mito_free_ for MDA-MB-231 cells, comparing cells from the population that invaded to those that did not invade into the collagen matrix. In all dot plots center lines are means with whiskers indicating the standard deviation in (a), (b) and (f). Statistical analyses concluded plotted data were not normally distributed with the Kolmogorov-Smirnov test, then performed pairwise comparisons with the Mann Whitney U test, where p-values were noted as: ns p≥0.05, *p<0.05, **p<0.005, ***p<0.001.

Inhibition of ROCK in 2D-cultured MDA-MB-231 cells resulted in cell elongation and branching, which was reflected in reduced circularity (Fig 6a). After adding Y-27632, there was a significant increase in fractional free cytoplasmic and mitochondrial NADH between untreated control values (30-45 minutes) and the average FPRM parameters (30-45 minutes) post treatment. Cyto_free_/Mito_free_ decreased after Y-27632 treatment. Due to the drastic elongated cell shapes that occur after Y-27632 treatment, cell body centroid tracking to measure migration distances was deemed to be inaccurate and migration distance was not included for comparison. In studies involving longer cell migration experiments, ROCK inhibition resulted in similar spindle-shaped morphological changes and an overall reduction in migration^53^.

In contrast, MDA-MB-231 cells in 3D collagen exhibited ovular shapes, and the addition of Y-27632 had no dramatic effect on cell morphology but there was a significant reduction in migration distance (Fig. 6b). Fractional free mitochondrial NADH increased with drug treatment and cytoplasmic NADH remained unchanged, resulting in a significant drop in Cyto_free_/Mito_free_. Both 2D and 3D changes show characteristics of more reduced NADH states and overall reduction of metabolic activities when exposed to Y-27632. Collectively, these data suggest that cells tune their bioenergetic needs in response to ROCK inhibition and culture dimensionality differentially affects to what extent shape changes may be involved.

### FPRM of vascular-cancer microfluidic co-culture system

Beyond monoculture 3D platforms, our group has previously characterized a 2-channel microfluidic platform in which one channel is seeded with human umbilical vein endothelial cells (HUVECs, endothelial cells (ECs)) and the other channel is seeded with cancer cells to study tumor cell migration in response to endothelial cell-secreted factors^28^ (Fig. 6c-d). Co-culturing MDA-MB-231 cells with ECs promoted a small population of cancer cells to migrate (“invading cells”) which exhibited differential pools of fractional free NADH as shown in Figure 6e-f. While no difference between fractional free mitochondrial NADH was observed, fractional free cytosolic NADH was significantly increased in invading cells compared to non-invading cells (Fig. 6e). This resulted in a significantly increased Cyto_free_/Mito_free_ in invading cells, suggesting a more metabolically active subtype (Fig. 6e). As characterized previously with metabolomics and computational flux balance analysis^28^, secreted factors from endothelial cells in this platform can induce tumor cell invasion into the surrounding collagen matrix by promoting ATP production, glycolysis, and oxidative metabolism, which validates our FPRM measurements. These examples highlight the versatility of FPRM in capturing the diverse spatiotemporal dynamics of single-cell migration in complex tissue culture platforms and provide insights into metabolic alterations that underpin cellular migratory behaviors.

## Discussion

We described an experimental pipeline to measure the metabolic status of individual cells using FPRM, an easy to implement imaging method based on the steady-state polarization state of two-photon excited NADH autofluorescence. FPRM requires an order of magnitude shorter acquisition time and does not induce the same levels of cellular stress as FLIM, which is commonly used to track NADH dynamics in cells and tissues. We applied FPRM to provide dynamic tracking of NADH signals with subcellular information during pharmacological and environmental manipulations, demonstrating the methods usefulness for monitoring metabolism. The use of ratiometric parameters such fractional free NADH in mitochondria and cytosol as well as Cyto_free_/Mito_free_ offer system independent readouts that are sensitive to metabolic changes and can be used to compare across different imaging setups. Furthermore, we coupled the dynamic changes of NAD(P)H signals with other parameters such as cell shape and migration speed in both 2D and 3D systems to establish the versatility of our imaging system in understanding cellular metabolism across different culture platforms at subcellular resolution.

In summary, we have developed and demonstrate a steady-state NADH emission polarization imaging method for monitoring cell metabolism that is faster and is less perturbing to cell metabolism then NADH-FLIM. FPRM approaches can be applied to advanced 3D culture systems to generate a more comprehensive understanding of the metabolic heterogeneity at subcellular and sub-minute resolution. We note that NAD(P)H polarization imaging would be affected by scattering when imaging deep into scattering samples, which may limit some experiments, however we achieved sufficient sensitivity to visualize changes in compartmental NADH status in cells embedded in scattering collagen hydrogels and expect engineered 3D culture platforms to be ideal settings to incorporate FPRM analysis. The simplicity and efficiency of FPRM is well-suited for high throughput studies of pharmacological activity of new drugs, as metabolic, morphological, and migration details can be sampled simultaneously with FPRM and heterogeneous responses are immediately clear. The power to dynamically track and correlate the metabolic and biophysical alterations of individual cancer cells will open doors to future biological discoveries of complex cellular behaviors in driving metastasis.

## Supporting information

Video listed in supplemental section 6

Video listed in supplemental section 6

## Acknowledgments

This work was supported by the Center on the Physics of Cancer Metabolism through Award Number 1U54CA210184-01 from the National Cancer Institute, R01GM132823, R01CA259195, and R01CA276392. Raman microscopy was performed through Cornell Center for Materials Research Shared Facilities which are supported through the NSF MRSEC program (DMR-1719875). All other imaging was performed through Cornell University’s Biotechnology Resource Center, with NIH S10OD018516 and NYSTEM C029155 funding.

## Author Contributions

L.L., J.C.C., M.L.T., and W.R.Z. performed experiments and data analysis. J.A.M.R.K facilitated L.L to operate Raman microscopy and performed data analysis. L.L, J.C.C, C.F, R.M and W.R.Z designed, supervised the experiments and wrote the paper. All authors were involved with editing the paper and discussion of results.

## RESOURCE AVAILABILITY

### Lead contact

Further information and requests for resources and reagents should be directed to and will be fulfilled by the lead contact, Warren R. Zipfel

### Materials availability

This study did not generate new unique reagents.

### Data and code availability

- Microscopy data reported in this paper will be shared by the lead contact upon request.
- ImageJ was used for image analysis and the basic workflow is presented in the image analysis section below, along with an example macro. The macros are modified depending on the experimental format and other macros are available upon request.
- Any additional information or code required to reanalyze the data reported in this paper is available from the lead contact upon request.

## EXPERIMENTAL MODEL AND STUDY PARTICIPANT DETAILS

### Imaging Media

Unless noted, all cell imaging was carried out in the following imaging media (buffer) we denote as IM: Seahorse XF DMEM with 5mM HEPES, without Phenol Red PN 103575-100 Agilent, supplemented with 10 mM glucose from Seahorse XF 1.0M glucose solution, PN 103577-100 Agilent with 1 mM glutamine added.

### Cell culture

MDA-MB-231 and MCF10A cells were obtained from ATCC. MCF10CA1a cells were obtained from Barbara Ann Karmanos Cancer Institute. Human adipose stem cells (ADSC) were purchased from Lonza (Basel, Switzerland). Adipose stromal cells (ASCs) were isolated from inguinal fat of 10 week-old female C57BL/6J (Jackson Laboratories) according to previously published protocols (Seo et al., 2015). MDA-MB-231 cells were cultured in Dulbecco’s Modified Eagle Medium (DMEM) supplemented with 100 U/mL Penicillin-Streptomycin (P/S) and 10% fetal bovine serum (FBS). ADSCs and ASCs were cultured in (1:1) Dulbecco’s Modified Eagle Medium:Nutrient Mixture F-12 (DMEM/F12) supplemented with 100 U/mL P/S and 10% FBS.

For adipogenic differentiation, ADSCs were induced with additional 1μM dexamethasone, 100μM indomethacin, 1μM insulin and 500μM 3-isobutyl-1-methylxanthine (IMBX) for 10 days. MCF10A and MCF10CA1a cells were cultured in DMEM/F12 media supplemented with 5% horse serum (HS), 10 μg/mL insulin, 0.5 μg/mL hydrocortisone, 100 ng/mL cholera toxin (Krackeler Scientific, NY, USA), 20 ng/mL epidermal growth factor (EGF), and 100 U/mL Penicillin/Streptomycin. All cell lines were maintained in incubators at 37°C with 5% CO_2_. Media was changed every 2 days until experiments. DMEM, DMEM/F12, P/S, FBS, HS, and EGF were purchased from Thermo Fisher Scientific (MA, USA). All other media supplements were purchased from Sigma-Aldrich (MO, USA), unless noted differently.

### Fluorescence Polarization Ratiometric Microscopy (FPRM) setup

Minor modifications were made to a Zeiss LSM 880 multiphoton/confocal microscope (Carl Zeiss GmbH, Oberkochen, Germany) to enable steady-state polarization imaging. Two-photon excitation was provided by an InSight DS femtosecond source (MKS/Spectra-Physics, Milpitas, CA) tuned to 730 nm emission to image NADH autofluorescence in cells. To ensure linear polarization of the excitation beam at the objective back aperture, the beam was routed through a Berek compensator variable waveplate (Model 5540, Newport Inc., Irvine, CA) placed after the NIR AOM. A Nichol Prism polarizer mounted on the microscope nosepiece and power meter was used to monitor the 730 nm laser power out of the microscope in order to determine the optimal Berek compensator settings for both X and Y polarizations (defined relative to the microscope stage). The two-photon generated epifluorescence was collected by the instrument’s four channel non-descanned detector (NDD) in which the fluorescence passed through a 680 nm short pass blocking filter (Chroma Technology Corp., Bellows Falls, VT) to further reduce any excitation light and then split by a polarizing beam splitter cube (PBS251 1” Polarizing Beamsplitter Cube, 420 – 680 nm, Thorlabs, Newton, NJ) housed in a modified Zeiss filter cube in the LSM 880 “BiG” detector unit. The separated P-polarized and S-polarized emissions passed through two matching 440/80 nm band pass filters (ET440/80m-2P, Chroma Technology Corp., Bellows Falls, VT) for detection by the gallium arsenide phosphide (GaAsP) photomultiplier tubes (PMTs) of the BiG unit (detectors “R3” and “R4” of the NDD unit).

The Berek compensator was set so that linearly polarized excitation light was parallel to the microscope stage’s X-axis (“X-polarized”), which directed the parallel polarization emission (P-polarized) into the “R4” PMT, while perpendicular emission (S-polarized) was detected by the “R3” PMT. P and S refer to the plane of incidence in relation to the polarizing beam splitter (P = parallel, S = senkrecht = perpendicular). On the three Zeiss LSMs we have available, we have found that the optical path to the external detectors (NDDs), as well as to a second two PMT unit located at the AiryScan port, have the same polarization reference frame as the excitation path and the emission polarization directions are preserved.

### Fluorescence Lifetime Imaging Microscopy (FLIM)

Two-photon fluorescence lifetime imaging microscopy was carried out using time-correlated single photon counting (TCSPC) module (SPC-830, Becker & Hickl Gmbh, Berlin, Germany) in a second PC with sync signals provided by the Zeiss LSM 880 system. Pulsed laser excitation at 730 nm (80 MHz repetition rate, ∼120 fs width at the sample) from the InSight DS femtosecond source coupled to the Zeiss LSM system. Two-photon generated epifluorescence entering the NDD unit passed through 440 nm band-pass filter (ET440/80m-2P, 316979, Chroma Technology Corp., Bellows Falls, VT) and detected by the “R4” PMT of the Zeiss BiG module operating in the photon counting mode. The excitation power was attenuated to laser power to keep the photon detection rate to less than 0.2% of the repetition rate, to avoid photon pile-up. The Insight’s 1040 nm output was reflected off a glass slide into a fast photodiode (FPD 310-FV, Menlo Systems Gmbh, Planegg, Germany) to generate the laser sync signal for the SPC 830 card. FLIM images were acquired using Becker & Hickl’s acquisition software (Spcm64.exe). Instrument response functions (IRF) were acquired by collecting TCSPC traces from Z-cut quartz generated SHG. Data was fit to a two exponential model using the SPCImage software from Becker & Hickl using the empirical IRF and the incomplete multi-exponential model provided in SPCImage. The amplitude-weighted mean lifetime was calculated as:

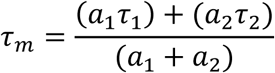

Where *τ*_1_ and *τ*_2_ are the lifetimes and *a*_1_ and *a*_2_ are the amplitudes of the two exponential decays of best fit.

### NADH in vitro binding assays

*In vitro* measurements of NADH polarization and fluorescence lifetime were made by titrating a 50 μM solution of NADH with the NADH-binding protein Glutamate Dehydrogenase (GDH) and fitting the measured response to the binding model described Supplementary Information section SI-2.2. Glutamate Dehydrogenase (GDH, MW) from bovine liver was purchased from Sigma-Aldrich (G7882, Type III, lyophilized powder, ≥20 units/mg protein) and resuspended to ∼ 23 μM in phosphate buffered saline (pH 7.4) with 1 mM EDTA. The concentration determination was based on A280 absorbance and a MW of 330 kDa (Sigma). Varying amounts of the stock GDH were added to 50 μM NADH in PBS. Hexamer GDH binds 7-8 NADH^1^ and the data in Figure 2 of the main text based on the protein concentration X 8.

### Sample preparation on 2D glass bottom dishes

In all 2D imaging experiments, unless otherwise mentioned, 35 mm glass bottom dishes (Matsunami, Osaka, Japan) were pre-coated with final concentration of 0.5 mg/mL rat tail collagen type I (Corning Inc., Corning, NY), then placed in incubator for 1 hour, and washed 3x with deionized water. Cells in culture around 80% confluence were trypsinized and the cell suspension was reseeded on the dishes at 10,000 cells per dish. After 24-hour incubation in the dish, the culture media was removed, IM added and 2D imaging experiments performed.

### Time course FPRM of unlabeled cells in 2D culture

MDA-MB-231 cells in IM were imaged on a Zeiss LSM 880 confocal microscope with 37°C incubation using the FPRM set up. Images were acquired at 5-minute interval for 1-4 hours depending on the experiment.

### Two-photon induced oxidative damage assays

#### Detection of protein glutathiolation

MDA-MB-231 cells were plated the day before the experiment on gridded coverslip bottom dishes (µ-Dish-35 mm, high Grid-500, Ibidi USA, Inc. Fitchburg, WI) and incubated for 1 hour in 1 mM BioGEE (Glutathione Ethyl Ester, Biotin Amide G36000, Thermo Fisher Scientific, Inc., Waltham, MA) in HEPES buffered DMEM/F12 phenol red free media preceding the experiment. Selected grid squares were irradiated using a Zeiss 20x/0.75 Fluar air immersion objective lens with ∼11 mW of 730 nm 120 fs pulsed light in the sample plane. Irradiation times were varied between 4 s, 60 s, or 240 s were carried out on separate grid dishes of MDA-MB-231cells. After illumination, the cells were permeabilized in Stabilization Buffer (SB: 4% PEG 8000, 1 mM EGTA, 10 mM PIPES pH 7.2) supplemented with 0.2% Triton X-100 for 30s, followed by 2x washes in SB (without Triton X-100). Cells were then stained with AlexaFluor-647 Streptavidin (S21374, Thermo Fisher Scientific Inc., Waltham, MA) for 20 min, then washed 3x in SB (without Triton X-100) and fixed in 4% pH adjusted Paraformaldehyde (PFA). Grid regions from the irradiation experiments were located and imaged on a Zeiss LSM 880 confocal microscope. Images of the irradiated grid regions and non-irradiated grid regions were analyzed with ImageJ. The Streptavidin-Alexa 647 staining was primarily localized to mitochondria and these regions were segmented out by thresholding, and histograms of the region averaged pixel values plotted.

### Cellular oxygen consumption measurement

A two-piece chamber with a 6 mm diameter round hole was milled from 6.35 mm thick plexiglass and #1.5 coverslip was glued to the bottom half using a UV-cured adhesive (NOA68, Thorlabs, Newton NJ). The bottom half of the chamber has two holes for stainless steel tubes that serve as the buffer input and output ports, and a third port for a phosphorescence-based oxygen probe (NeoFox Oxygen probe, Ocean Optics, Orlando, FL, USA). An O-ring groove was milled into the top piece of the chamber which is screwed to the bottom half to create a seal. The imaging chamber volume was ∼0.2 ml. MCF10a cells were plated in the well with the coverslip bottom and after 24 hours the culture media was exchanged for an ambient oxygenated IM buffer (∼210 μmol/L O_2_ at 37C). The imaging chamber was placed upside down on an upright Zeiss LSM 880 confocal/multiphoton system and oxygenated IM buffer was perfused through the imaging chamber. When perfusion stops a decrease in the O_2_ is seen due to respiration, with the rate of change being the relative respiration rate of the cells. The perfusion was turned on and off several times as the electrode response was recorded. The decrease in oxygen after perfusion was stopped was fit to an exponential decay model to obtain to a measure of the oxygen uptake by the cells. To assess changes due to two-photon excitation, the respiration rate was also recorded after 200 seconds of scanning with ∼10 mW of 730 nm femtosecond excitation delivered through a 20x/1.0 NA objective lens. Respiration data could not be recorded during the irradiation period due to oxygen electrode artifacts from the NIR laser illumination.)

### Live cell organelle staining

#### Co-staining of mitochondria for correlation with 440nm autofluorescence

Cells prepared on glass bottom dishes were retrieved from an incubator, and stained with 1 µM MitoTracker Red CMXRos (Thermo Fisher Scientific Inc., Waltham, MA) in the respective cell line’s growth medium for 10 min. All stained cells were rinsed 3x with fresh phenol red-free, HEPES-buffered DMEM/F12 media to remove excess dye before imaging.

#### Co-staining of neutral lipid for correlation with 440nm autofluorescence

Cells prepared on glass bottom dishes were retrieved from an incubator, and stained with 1 µM Nile Red (Thermo Fisher Scientific Inc., Waltham, MA) in the respective cell line’s growth medium for 10 min. All stained cells were rinsed 3x with fresh phenol red-free, HEPES-buffered DMEM/F12 media to remove excess dye before imaging.

#### Imaging

Images were acquired on LSM 880 microscope with both FPRM channels and the organelle channel simultaneously for co-localization comparison. A 560 nm short pass dichroic was inserted in the Zeiss NDD external detector to direct emissions above 560 nm into the second PMT of the four-channel unit, passing the shorter wavelengths on to the two PMT BiG module which housed the polarizing beam cube.

### Fluorescence emission spectra

Intracellular fluorescence emission spectra from 415 to 690 nm were acquired using the LSM 880’s 32-channel spectral detector set for a 9 nm bandwidth per channel. Two-photon excitation was at 730 nm delivered through a 63x oil/1.4 oil immersion objective.

### Raman

Mapping was done using a confocal Raman microscope (alpha300R, WITec, Ulm, Germany) with a Diode-Pumped Solid-State laser (532nm) excitation source. The Raman scattering cross section was analyzed with a UHTS 600 spectrometer (WITec, Ulm Germany) equipped with 300 l/mm grating and high efficiency (QE>90%), back illuminated, CCD camera (look up model number, Andor, UK). Laser power was measured with a power meter (S130C, Thor Labs) and set to 63 mW. For each cell type, a glass bottom dish containing fixed cells submerged in phosphate buffer was placed on a motorized stage for sample positioning and imaged using a 63X water immersion objective (Leica, NA = 0.9). The theoretical lateral resolution was ∼0.61 λ/NA = 361 nm. The step size was 1 μm with an integration time of 0.3 s per step.

### Time course FPRM of 2D cell culture during metabolic inhibition

MDA-MB-231 cells prepared on glass bottom dishes were retrieved from an incubator, and the media was replaced with 1 mL of IM and relocated to the FPRM microscope’s heated stage incubator. Tiled or multi-position FPRM images were acquired at 5 min intervals over a time-course experiment. For each sample, three FPRM images were taken at the unstressed initial steady-state condition, and immediately after the third timepoint (designated t=0 min), 1 mL of one of the metabolic inhibitors in 37°C IM was added at 2x concentration to the sample. For 2-deoxy-G-glucose (2DG), the final concentration was 50 µM; for potassium cyanide (KCN), the final concentration was 5 µM; for carbonyl cyanide 4-(trifluoromethoxy)phenylhydrazone (FCCP), the final concentration was 1 µM and for AZD7545, the final concentration was 5 µM. The media addition mitigated physical contact with the microscope, and the time course imaging experiment was continued until 40-60 minutes of imaging had elapsed.

### Hypoxia/Normoxia experiment

35 mm glass bottom dishes (D35-10-1.5-N, Cellvis, Mountain View, CA) were pre-coated in 50 µg/mL Collagen type I purified from rat tail (354249, Corning Inc., Corning, NY) by incubation at 37°C for 1 hr. Glass surfaces were then washed 3x in phosphate buffered saline (PBS) and 1x in MDA-MB-231 culture medium. MDA-MB-231 cells in culture around 80% confluence were trypsinized to rinse and count, and the cell suspension was reseeded on the collagen-coated dishes at 100,000 cells per dish. All samples were set in an incubator with standard incubation conditions. After 24 hr. of incubation, samples were split into two groups: Normoxia and Hypoxia. The normoxia treatment group continued to be incubated in 37°C, ambient O_2_, and 5% CO_2_ culture settings, while the hypoxia treatment group was moved to a similar incubator with 37°C, 1% O_2_, and 5% CO_2_. After 3 days of incubation, samples were washed 1x in 1 mL of 37°C IM, then left with fresh 1 mL of IM and moved to a pre-heated stage incubator for FPRM imaging with minimal delay (as the stage incubator maintained ambient O_2_).

### Glucose vs Glucose-Free experiment

35 mm glass bottom dishes (D35-10-1.5-N, Cellvis, Mountain View, CA) were pre-coated in 50 μg/mL Collagen type I purified from rat tail (354249, Corning Inc., Corning, NY) by incubation at 37°C for 1 hr. Glass surfaces were then washed 3x in phosphate buffered saline (PBS) and 1x in MDA-MB-231 culture medium. MDA-MB-231 cells in culture around 80% confluence were trypsinized to rinse and count, and the cell suspension was reseeded on the collagen-coated dishes at 100,000 cells per dish. All samples were set in an incubator with standard incubation conditions. After 24 hr. of incubation, samples were split into two groups: some with glucose and some without glucose. Samples were washed 1x in 1 mL of 37°C Imaging Medium (IM, as described above), then left with fresh 1 mL of IM (0 mM Glucose for the glucose-free treatment; 10 mM glucose for the glucose treatment group) and moved to a pre-heated stage incubator for FPRM imaging after 30 minutes to acclimate to the media adjustment. Single timepoint tile-scan FPRM images were acquired from each sample and analyzed using the image analysis pipeline described below.

### Time course FPRM of unstressed and ROCK inhibition in 2D cell culture

Tiled FPRM images were acquired at 5 min intervals over a time-course experiment. For each sample, cells in 1 ml media were imaged at the unstressed initial steady-state condition for 60 minutes. Immediately after 60 minutes, 1 mL of Y27632 was added at 2x concentration to the sample with the final concentration of 30µM, which inhibits rho-associated protein kinase (ROCK) signaling. The inhibition time course imaging experiment continued until another 60 minutes of imaging had elapsed. For calculation, control values were calculated from 30-45 minutes in unstressed conditions, and Y27632 inhibition values were calculated from 30-45 minutes after adding Y27632.

### Sample preparation of 3D collagen-embedded cells

Collagen microwells were prepared as previously described (Ling et al., 2020). In brief, polydimethylsiloxane (PDMS, Slygard 184 Silicone Elastomer kit, Dow Corning, Midland, MI) was cast into a 10cm petri dish with a thickness of 1 mm. A ring-shaped PDMS well was stamped from the cured PDMS slab by punching concentric 8 mm and 10 mm diameter disks with biopsy punches (Integra LifeSciences, Princeton, NJ). Individual PDMS rings were plasma treated (Harrick Plasma) and covalently bonded to individual glass bottom dishes (Matsunami, Osaka, Japan). The inner regions of PDMS rings were treated with 1% poly-ethylenimine (Sigma-Aldrich, St. Louis, MO) and 0.1% glutaraldehyde (Thermo Fisher Scientific Inc, Waltham, MA) to promote collagen attachment to the wells. Rat tail collagen I (Corning Inc., Corning, NY) was reconstituted and neutralized with sodium hydroxide and 10x concentrated DMEM/F12 media to a final concentration of 6 mg/mL before mixing with 800,000 cells/ml. The collagen gel mixtures were incubated at 4°C for 15 min, 20°C for 15 min, then 37°C for 15 min to promote the formation of collagen fibers. Each gel mixture was then immersed with media and cultured at 37°C for 1 day before switching to imaging media (IM: Seahorse XF DMEM with HEPES and supplements).

### Time course FPRM of 3D cell culture during ROCK inhibition

Cells were imaged at steady state (in microscope stage incubator) at 15 min intervals for 1 hr. before adding Y-27632 (R&D Systems Inc., Minneapolis, MN) at 30 µM final concentration. Imaging continued for 2 hr. after application of the inhibitor. For calculation, control values were calculated from 30-45 minutes in unstressed conditions, and Y27632 inhibition values were calculated from 135-150 minutes after adding Y27632.

### Sample preparation of 3D collagen luminal microfluidic device

#### Preparation of the device

Microfluidic devices were generated by pouring a degassed polydimethylsiloxane (PDMS) solution with a 1:10 ratio of curing agent and elastomer onto a silicon master and cured overnight at 60°C. PDMS devices were subsequently removed from mold and media wells were cut out using a square biopsy punch. PDMS devices were then cleaned using O_2_ plasma for 5 minutes before bonding to #1 cover glass. To facilitate collagen hydrogel attachment, assembled devices were treated with 1% v/v polyethylenimine in water and 0.1% v/v glutaraldehyde in water for 10 and 30 min respectively before washing with water overnight. Devices were subsequently UV sterilized for 10 minutes before the insertion of 300 μm diameter acupuncture needles coated with 1% BSA in PBS. A neutralized 2.5 mg/ml solution of rat tail collagen type I (Corning) was injected into devices, polymerized at 4°C for 30 min followed by a final incubation at 37°C for 30 minutes. Needles were removed to generate hollow microchannels within the collagen hydrogel which were subsequently rinsed with EGM-2 for 24 hrs.

#### Culturing cells in the luminal microfluidic device

For co-culture experiments, a suspension of 2 x 10^6^ per ml human umbilical vein endothelial cells (HUVECs) was prepared and 10 μL of this suspension was injected into one channel of the device. Device was flipped upside down and incubated at 37 °C for 10 min to facilitate attachment of HUVECs to upper side of channel. A second seeding was then performed after flipping devices right-side up, incubating for another 10 min. Media was replaced with fresh EGM-2, and devices were incubated at 37°C on a rocker set to a 15°C tilt angle and 2 RPM for 48 hrs.

Next, a cell suspension of 2 x 10^6^ MDA-MB-231 per ml was prepared in EGM-2 and 10 μL of this suspension was injected into one port of the remaining channel, followed by another 10 μL into the opposite port of the channel. Devices were incubated for 30 min at 37°C before changing media. High glucose DMEM supplemented with 1% FBS, 1% P/S (low serum DMEM) was added to the channel containing MDA-MB-231, while fully supplemented EGM-2 was added to the channel containing HUVECs. Devices were placed back on the rocker in the incubator and cultured for 3–5 days. Media was replaced every 24 hrs.

#### Immunofluorescence staining

To fix microfluidic devices, media was removed from all reservoirs and replaced with 4% PFA, incubating for 20 min temperature. After fixation, devices were rinsed with PBS and subsequently permeabilized with 0.1% v/v Triton X-100 in PBS for 15 minutes. Devices were then blocked with 2% w/v BSA in PBS (blocking buffer) for 1 hr at room temperature. Primary staining was performed by incubating with anti-CD31 (Sigma, 1:100, clone WM-59) diluted in blocking buffer overnight at 4°C. After washing with PBS for 24 hrs, secondary staining was performed by incubating with goat anti-mouse Alexafluor-647 (Thermo Fisher Scientific), DAPI, and Alexafluor-568 Phalloidin for 2 hrs. at room temperature. Stained cultures were washed with PBS overnight at 4°C before imaging.

#### Confocal fluorescence imaging of stained sample

To obtain fluorescence images of fixed microfluidic devices, a 10×/0.45 C-Aprochromat objective with a 0.6 optical zoom was used to generate maximum projected z-stack tile-scans. For FPRM imaging, fresh media was added to devices and PDMS slabs were placed over media reservoirs to prevent evaporation due to a heated stage. A 32x/0.85 C-Achroplan objective was used, obtaining images of MDA-MB-231 that remained within the channel (non-invaded) and those at the invasive front within the collagen hydrogel (invaded).

### NADH *in vitro* FPRM data analysis

See Supplemental Information SI-1 for theoretical discussion and fitting results.

### NADH *in vitro* FLIM data analysis

Using B&H software, a grayscale image representing the number of counted photons per pixel (“Intensity Image”) and a correlated image representing the computed weighted mean fluorescence lifetime per pixel (“Color-Coded Image”) were exported and plotted for analysis.

### Raman Microscopy data analysis

Hyperspectral data (image-graph objects) were analyzed using WITec Project 5.1 software. Data were first spectrally cropped from 328-3770 cm-1. They were then background subtracted using the ‘Graph Background Subtraction’ dialog (Tab: Shape, Shape Size: 500, Keep Border Peaks: unchecked). Peak area images were generated using sum filters in the ‘Filter Viewer’. Lipids were captured using the characteristic peak at 2850 cm-1. The range was 2825-2855 cm-1 and the background was defined as a constant, anchored to the left bound (set to the average intensity of four points nearest 2825 cm-1 and zero at the 2855 cm-1). The nuclear/cytoplasm signature (DNA and proteins) was integrated from 2928-2988 cm-1, where the background was defined as linear and anchored at both bounds as the average of intensity of four points nearest each range limit. The combination image shown in Fig 3d was generated using the Image Combination Dialog. Auto thresholding was used for the upper value of the images. The bottom value was set to zero (Color Mixing: unchecked). To generate the spectra shown in Fig.3d and SI-4, the top 5% of positive valued pixels from the peak area images were extracted and averaged. First, peak area images were examined in the ‘Image Histogram and Statistics’ dialog (full range of pixels). In the Histogram tab, the lower value (Range Start) was set to zero and these (negative) pixels were excluded in the Draw Field Mask tab (Clear Out of Range Pixels). Next, in the Statistics tab the 95% Level value was identified and used as the new lower limit and all pixels below this were similarly excluded. The remaining pixels were extracted as a mask. The mask was further multiplied by a manually made cosmic ray mask to remove cosmic rays. These remaining pixels were averaged to generate the spectra. For MCF10CA1a. the number of pixels averaged for the DNA + cytoplasm-rich spectrum was 107 and for the lipid-rich spectrum was 79. For MCF10A the number of pixels averaged the lipid-rich spectrum was 68.

### Analysis for NAD(P)H FPRM imaging of cells on 2D substrates

All image data is acquired as 16-bit data. 2D image analysis was carried out using a series of ImageJ macros. Depending on the experimental imaging setup/protocol (time series, multiple fields of view stored as a single CZI, etc.) the macros are modified as needed. Below is an outline the main image processing steps used to extract the compartmental pixel values used to calculate the fractional free NADH and an example ImageJ/FIJI macro.

1. The first macro creates a series of TIF images from the Zeiss CZI files. These are typically multi field-of-view XYT files (Tiled region and a series of selected regions across the dish). The two channels (S, perpendicular and P, parallel polarizations) are separated and saved as ImageJ TIFs, and the sum of the S and P channels is calculated as a 32-bit intensity image, reduced to 16-bit data and saved as an ImageJ TIFF. The intensity file is used to create the binary masks. Tiled images (taken with 10% overlap) were stitched using the Zeiss Zen software before channel separation and conversion to ImageJ TIFs.
2. The summed intensity image is used to create threshold based binary masks for the whole cell outline and mitochondria, which are saved as 8-bit TIFs. The mitochondrial mask is inverted and divided by 255, and the cell mask multiplied by the normalized mitochondrial mask to create the cytoplasm mask (see Figure 2 of the main text).
3. A third macro opens the binary mask files and divides mitochondrial and cytoplasm masks by 255. For analysis of the mitochondria and cytoplasmic regions, the S and P images are divided by the normalized mitochondrial mask and cytoplasmic masks using the ImageJ “Image Calculator” creating floating point images (denoted as “**`cytoS`**” and “**`cytoP`**” in the code below) where all pixels outside of the mask are marked as NaN (“not-a-number”). At the start of the macro the command ***run("Misc…", "divide=NaN run");*** should be used to ensure that both x/0 and 0/0 are set equal to NaN’s when encountered. Using the cell mask Regions of Interest (ROIs) can be selected of individual cells (or cell clusters) and cytoplasmic and mitochondrial masks applied to find the mean S and P intensity values for those regions.

The **`Smean`** and **`Pmean`** values in the macro below are used to calculate the fractional free NADH in mitochondria (Mito_free_) and cytosol (Cyto_free_) using previously measured S/P_free_ and S/P_vitrified_ values and equation 1 from the main text. The normalization values S/P_free_ (NADH in PBS, typically ∼20 μM) and S/P_vitrified_ (Blue plastic slide from Chroma Technology, Bellows Falls, VT) should be measured at the start of each set of experiments, especially if any system components have been changed (e.g. different objective lens). The results are saved within the macros as ascii text files for plotting and further analysis.

Example macro:

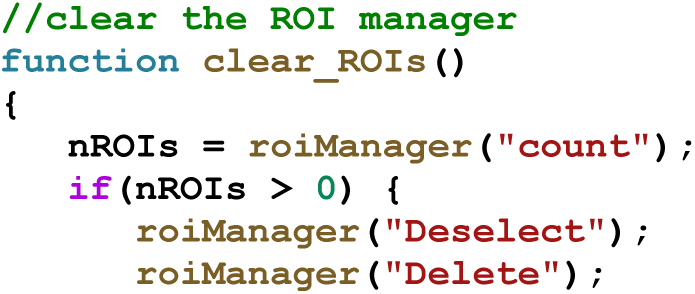
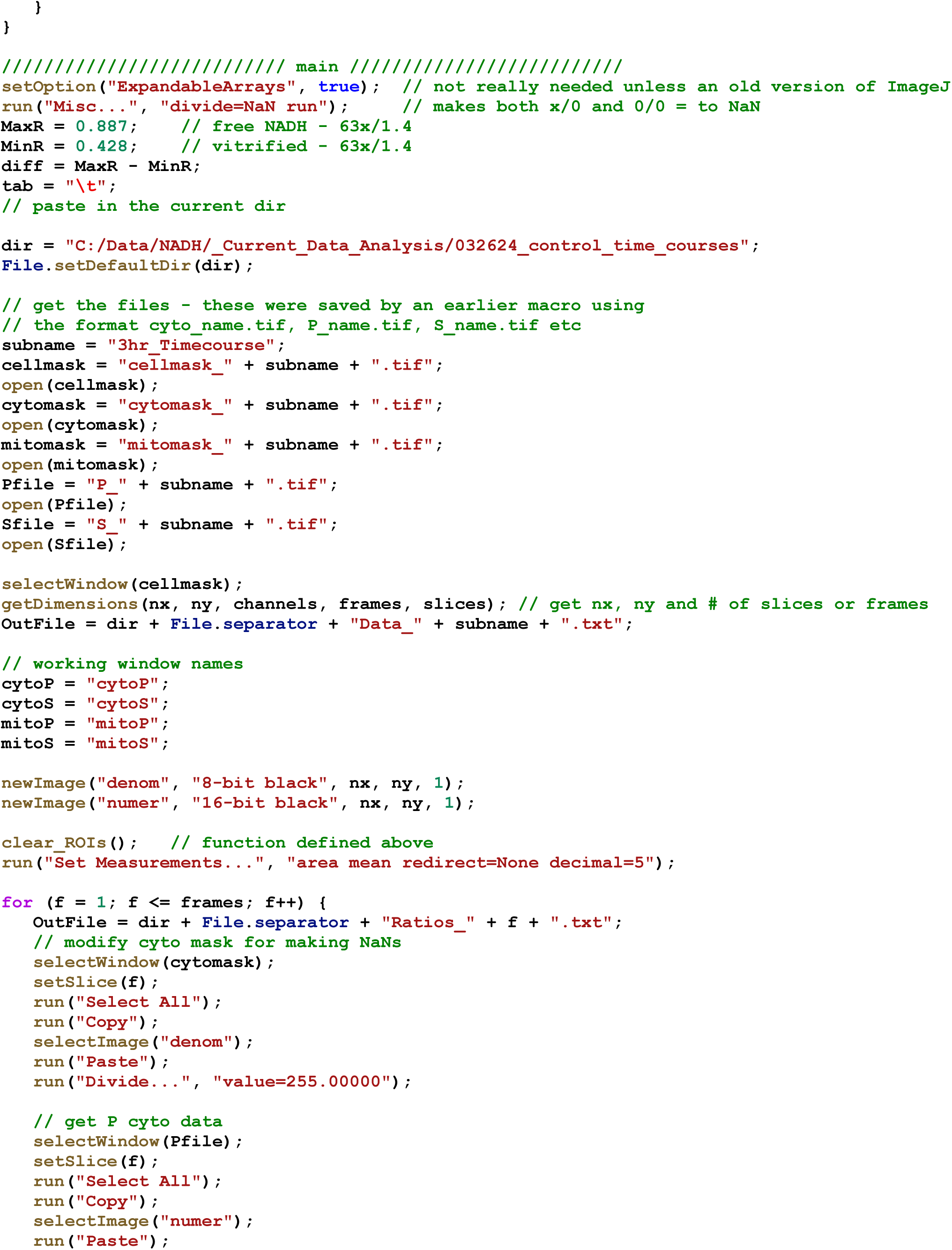
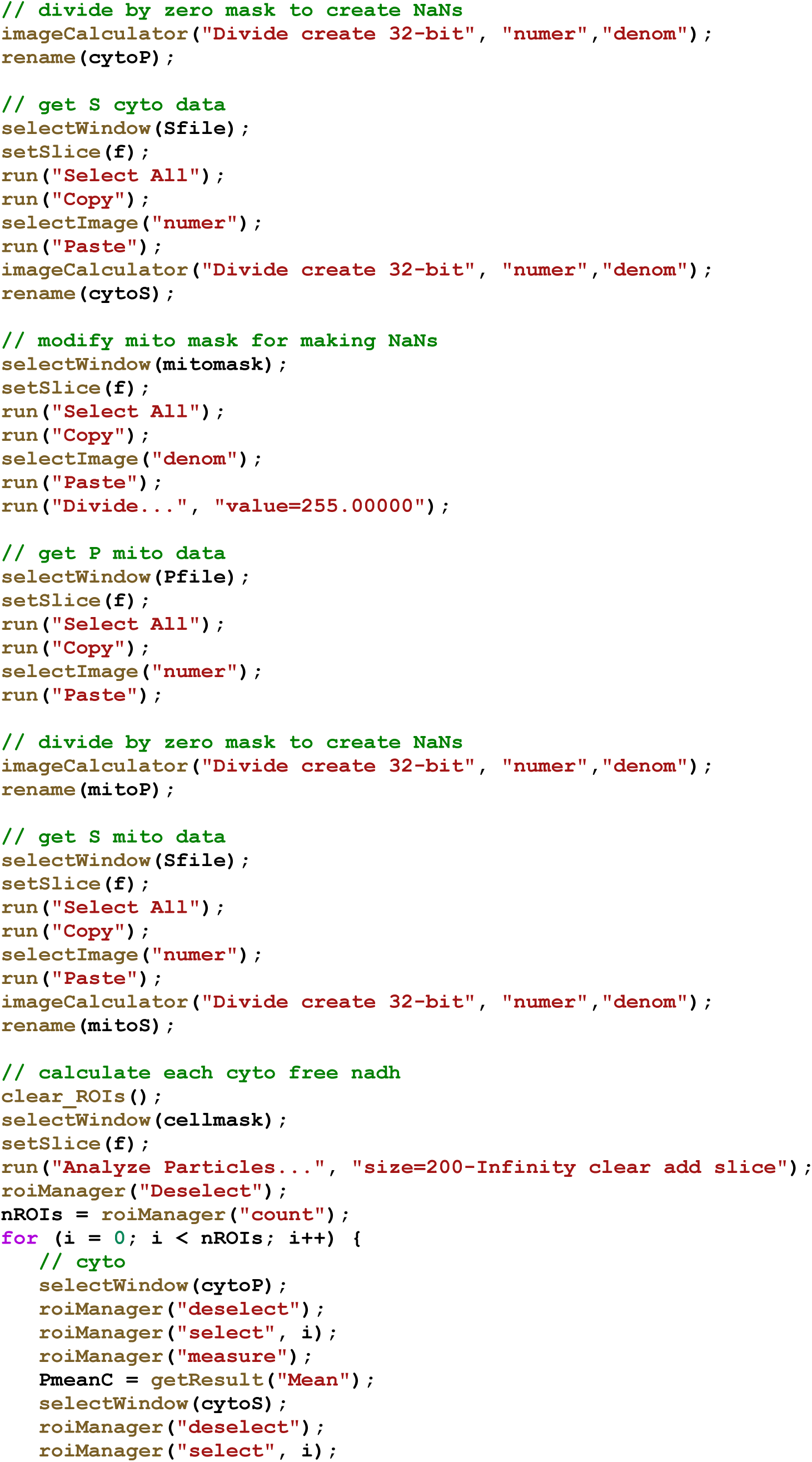
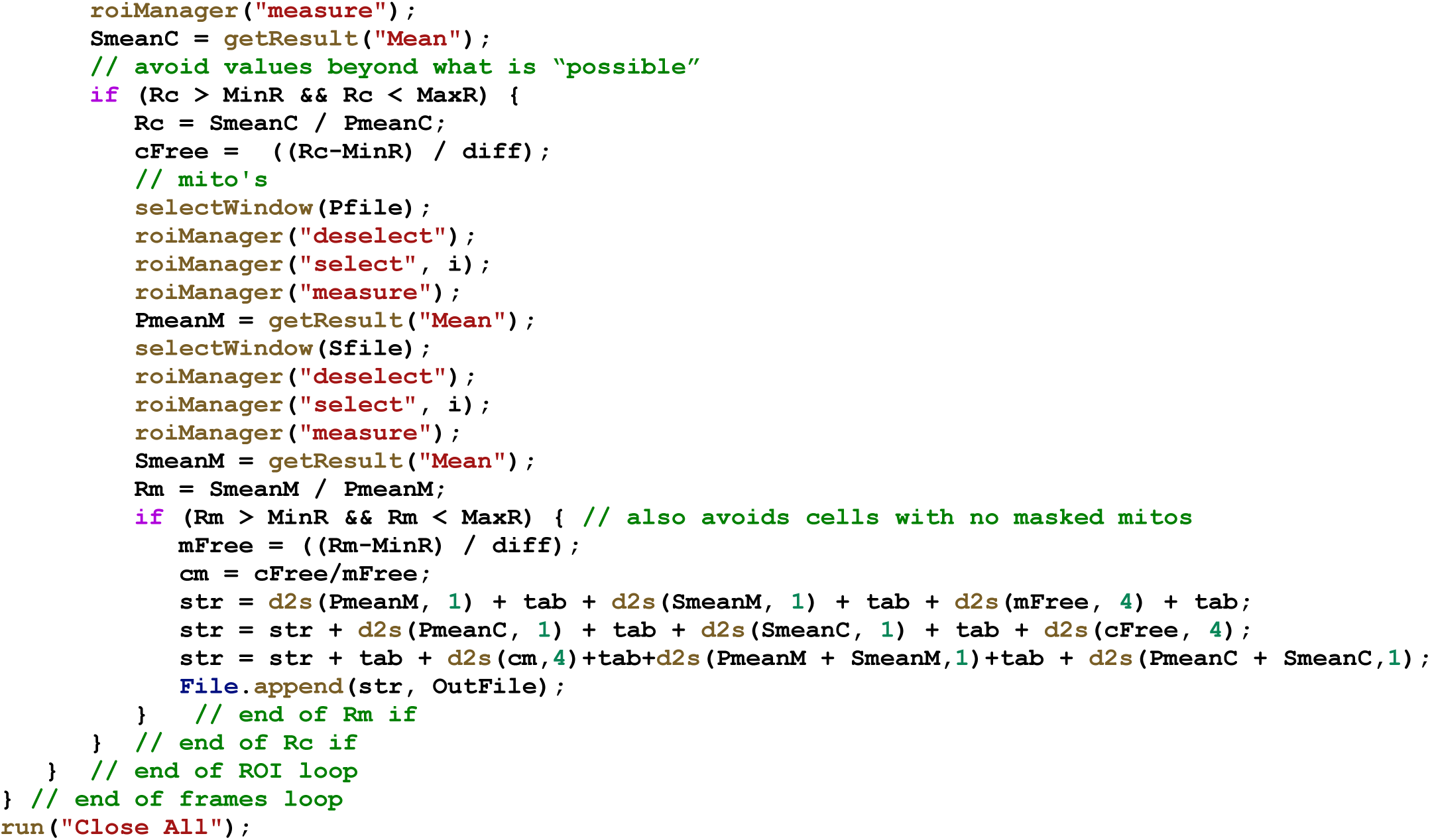

#### Other image parameters

Shape descriptor (circularity) and centroid (x, y information) were calculated from the cell masks. Results are saved by the macros as ascii text files for plotting and further analysis.

### Analysis for NAD(P)H FPRM imaging of cells in 3D engineered matrices

Confocal z-stacks were analyzed using custom ImageJ macros to quantify 3D measurements. Briefly, regions of interest (ROI) for cells were defined using maximum intensity projections of channels of interest. Afterwards, slices of the z-stack restricted to the selected ROI and only including one cell body at each time point were manually saved as individual Z-stacks named according to cell index. The central five Z-slices were used per cell for calculations. Intensities were measured per slice like 2D analysis and the mean values acquired from the 5 slices were calculated per cell. The centroid (x, y, z) was calculated using “3D Objects Counter” in ImageJ from the individual Z-stack for each cell, with threshold of intensity of minimum of 3000 and maximum of 87885. The migration distance was calculated as:

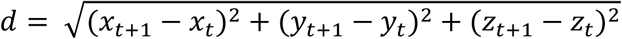

### Statistical analysis

Detailed experimental set ups were described in previous methods section and figure legends. Microsoft Excel, OriginLab, GraphPad Prism, and JMP were used for statistical analysis. Specific details of each test used were described in each figure caption.

### Illustrations

Data was plotted in OriginPro 2024 (OriginLab) and scientific illustrations were created using Adobe Illustrator v29.5.1.

## KEY RESOURCES TABLE

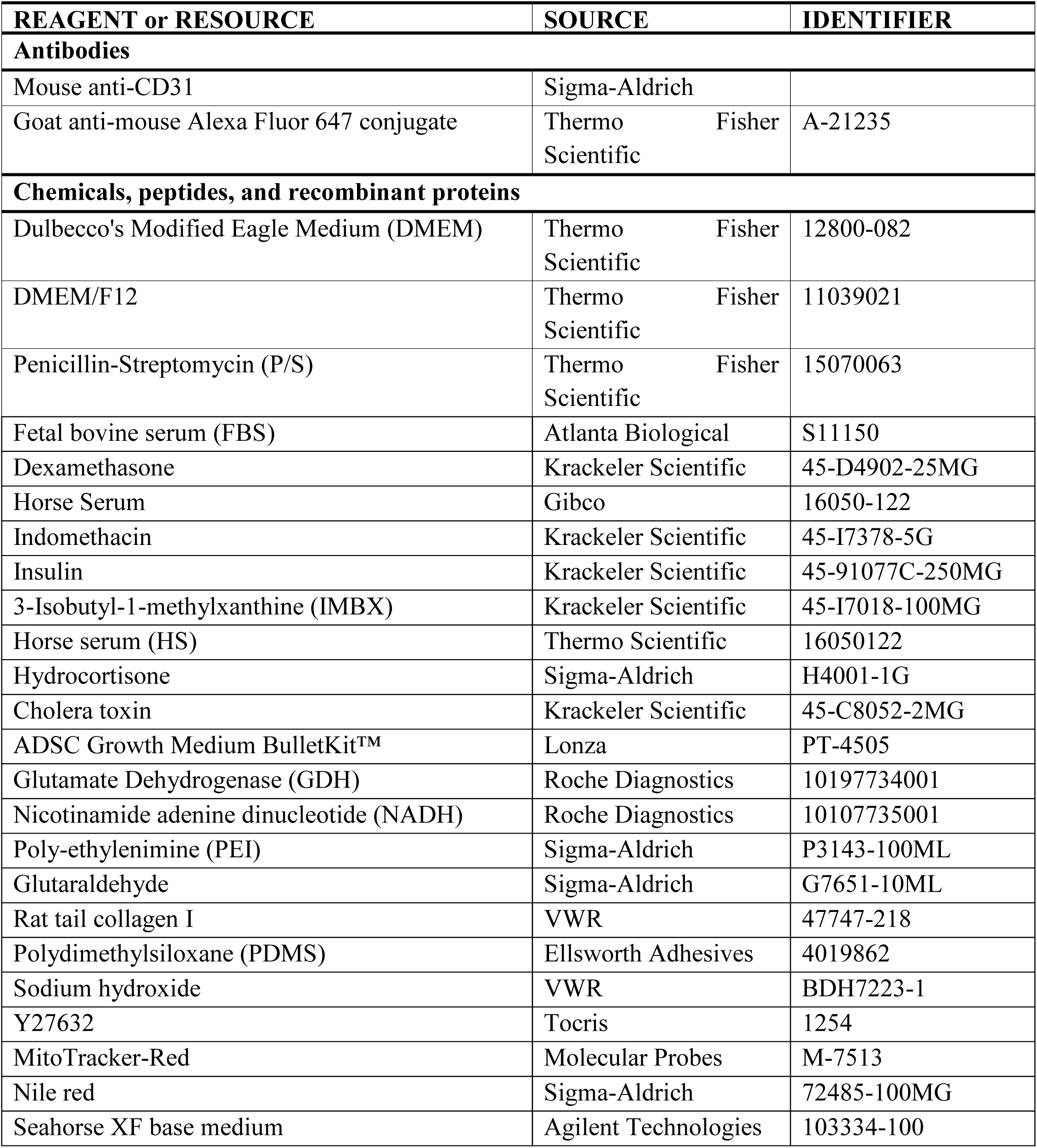
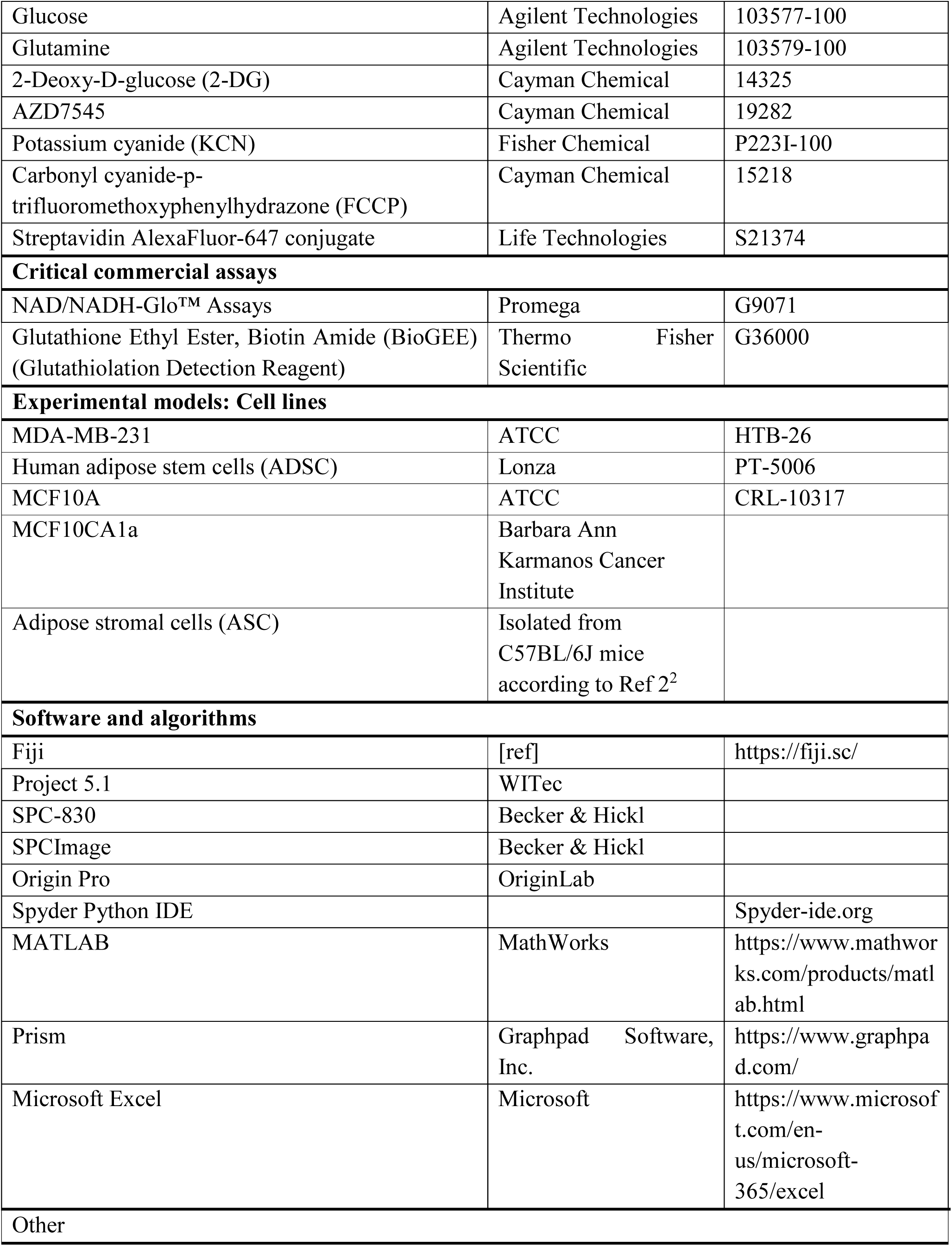
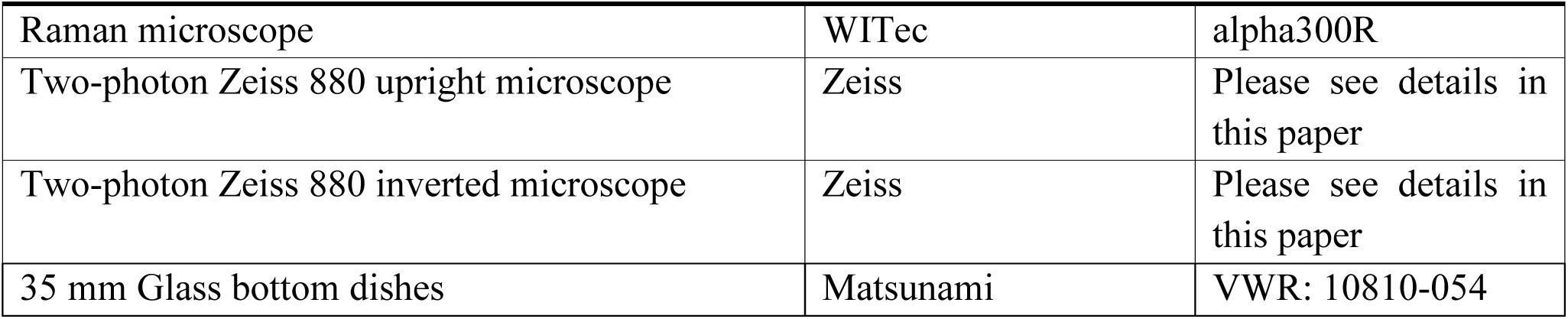

1 Shafer, J. A. *et al.* Binding of Reduced Cofactor to Glutamate Dehydrogenase. *European Journal of Biochemistry* **31**, 166-171, doi:https://doi.org/10.1111/j.1432-1033.1972.tb02515.x (1972).

2 Ling, L. *et al.* Obesity-associated Adipose Stromal Cells Promote Breast Cancer Invasion Through Direct Cell Contact and ECM Remodeling. *Adv Funct Mater* **30**, doi:10.1002/adfm.201910650 (2020).

## Supplementary Information

### Section SI-1. Two-photon NAD(P)H excitation and fluorescence emission rates

The two-photon action cross-sections (two-photon absorption cross-section x the quantum yield) of free and protein bound NADH (Fig. SI-1a) are typically 2 to 3 orders of magnitude lower than standard fluorescent dyes. NADH only very rarely emits a photon after excitation with a tiny quantum yield (0.02 for unbound to ∼0.08 when bound^1^) which significantly limits the acquisition rate. The low quantum yield means that > 90% of the absorbed energy is going to photophysical processes other than fluorescence, potentially causing photodamage. The number of photons collected during our typical FPRM 6 μs pixel time (4 frame average) is ∼10 to 40 (∼1 to 7 photons/μs), with 10-20 photons/pixel typical with excitation powers below ∼10 mW for viability reasons.

**Figure SI-1a and b.**
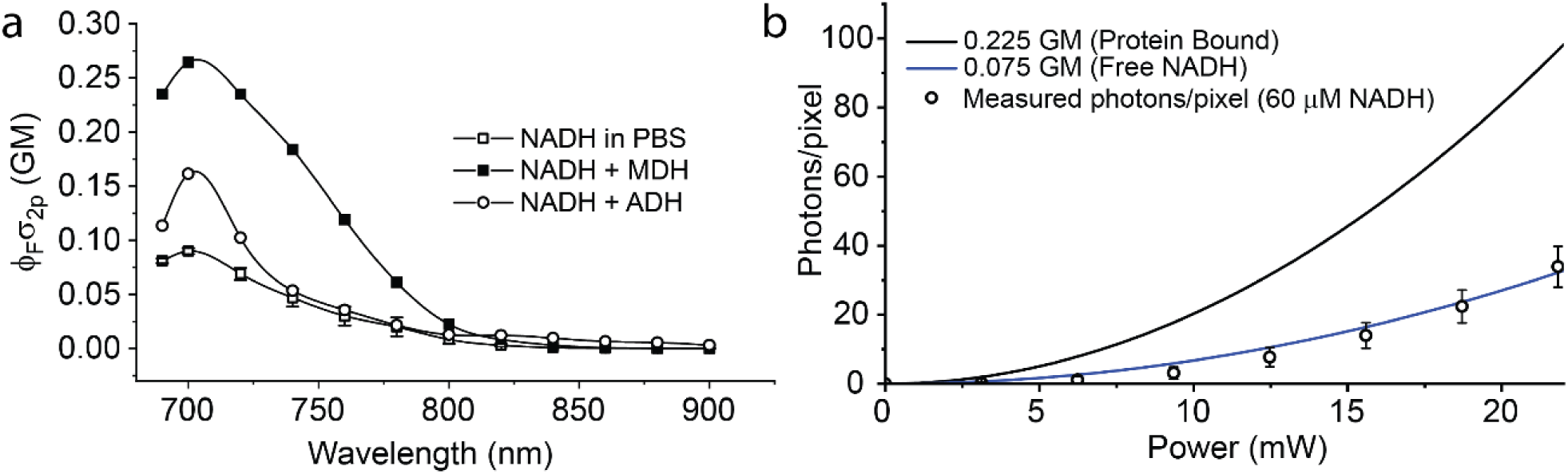
NAD(P)H photophysics. **(a)** Two-photon action spectra of free NADH in solution and NADH bound to two different NADH-binding enzymes. 2P action cross-sections were measured as described in Ref 2^2^. For protein bound measurements, 50 μM NADH was titrated with protein (MDH, ADH) and fluorescence monitored to find the protein concentration at which the emission intensity saturated. Solutions at this concentration without NADH were used to correct for any background two-photon excited signal from the protein. (b) Based on the two-photon action cross-sections, nonlinear excitation parameters (laser power, pulse width etc.), numerical aperture and collection efficiency, and the expected number of NADH fluorescence photons detected per second can be estimated. Lines in (b) are calculated using the typical parameters used in FPRM of NADH (Appendix SI-1) and assuming 60 μM NADH. Circles along the NADH in solution line are measurements made in photon counting mode on a Zeiss LSM-880 (mean ± sd photons per pixel) for 60 μM NADH in PBS.

The low action cross-section is also the fundamental limiting factor for NAD(P)H FLIM. With standard Time Correlated Single Photon Counting (TCSPC) the detection rate must be significantly less than 1 photon per laser pulse to avoid statistical sampling errors, since only one photon can be processed per Time to Amplitude Converter (TAC) cycle. Although some newer TCSPC systems can process multiple photons per laser pulse and deadtimes have also been reduced (the SPC-830 TCSP card used in this work has a ∼100 ns deadtime between detected photons), if an accurate two-component fit is needed it would require ∼4 x10^5^ total photons per decay trace^3^ for each pixel. In raster scanning microscopy a pixel is sampled every 0.5 to 1 second (assuming typical frame rates), which is why NADH-FLIM requires long FLIM frame acquisition times. Less than one photon is detected per laser pulse (∼0.01 to 0.5 photons per pulse on average, based on the photophysics described above) and even with a zero deadtime TCSPC card, it will always take minutes to collect a high temporal resolution FLIM image.

### Section SI-2. Polarization based imaging as implemented in FPRM

#### 2.1 FPRM normalization

Polarization methods, as implemented on a fluorescence spectrometer, typically use linearly polarized plane wave excitation and a single detector with a rotatable polarizer to sequentially sample the perpendicular and parallel emissions. Fluorescence polarization data taken in this way is most commonly calculated as fluorescence anisotropy (*r*) defined as:

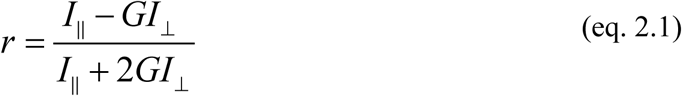

Fluorescence anisotropy values are reproducible across spectroscopy instruments because a system dependent correction factor (“G” – the grating factor) is used to compensate for the polarization-dependent emission grating reflectivity. Although this approach has also be used for implementing an imaging anisotropy mode on a laser scanning microscope as in Smith et al^4^, it is not required. FPRM uses two separate PMTs with differing intrinsic gains, as well as instrument-dependent optical factors that affect the polarization. such as objective birefringence, NA and optical path differences. In FPRM we define a factor α which takes PMT gain differences, the extent of axial polarization introduced by the high numerical aperture focus (Fig. SI-2.3.1), and any polarization effects from the optics used (e.g. objective birefringence or from emission path dichroics and filters) into account. We normalize the S/P ratio to a value bounded by the S/P of free NADH in solution and the S/P ratio from a vitrified sample (blue plastic slide or NADH in a polymer) canceling α and providing an estimate of the factional free NADH:

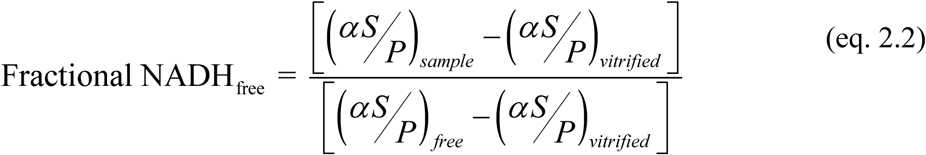

We find that this corrects for the factors listed above and makes the method instrument independent as shown in section SI-2.3. This method slightly overestimates the free NADH at the low end because fully bound NADH in a biological milieu never reaches the vitrified polarization value. However, this treatment enables an easy normalization that is robust day-to-day (see section 2.3).

##### Steady State polarization ratio and fluorescence lifetime

The S/P ratio depends both on the rotational time of the molecule and the radiative lifetime of the fluorophore (Perrin equation):

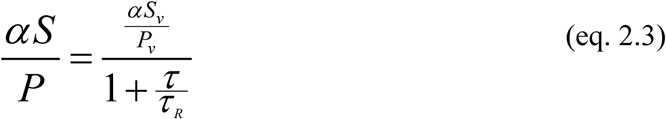

Where S_v_ and P_v_ are the intensities for a vitrified sample or “frozen” sample (e.g. a fluorophore immobilized in a polymer), τ is the fluorescence lifetime and τ_R_ is the characteristic rotational time of the molecule. Although the NADH lifetime increases when bound, the rotational time of NADH increases significantly more, accounting for the S/P values we correlate with cell metabolic state. The rotational time of NADH in solution is around 0.3 - 0.4 ns (∼equal to the decay time of free NADH). Although τ_R_ is slightly longer in cytosol due to the higher viscosity, when protein-bound it gets much longer than the fluorescence lifetime increases and 1 + τ/τ_R_→1. In addition to having a more polarized emission, bound NADH also becomes brighter, as shown in Figure SI-1a. This adds more complexity to both steady-state polarization measurements and FLIM, since the bound form is more heavily weighted and the quantum yield increase depends on the binding partners (which vary by compartment, and possibly cell type).

#### 2.2 Kinetic model of PS vs enzyme binding

Figure 2a in the main text shows a fit (red line) of the P/S ratio to a model based on the fractions of free and bound NADH and the quadratic binding equation:

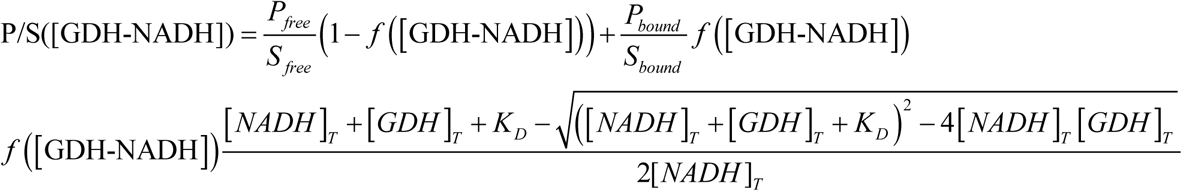

The equation above is in terms of P/S to model the data shown in Figure 2b in the main text. Fitting the kinetics (P/S vs a varying concentration of glutamate dehydrogenase NADH binding sites: [GDH]_T_) using standard nonlinear least squares (Levenberg-Marquardt method) to the equation above results in a K_D_ value of 26 μM for NADH binding. *f*[GDH-NADH] is the steady-state fraction of NADH bound to GDH; [NADH]_T_ and [GDH]_T_ are the total steady-state concentrations; [NADH]_T_ is held constant at 50 μM and [GDH]_T_ varies. (P/S)_free_ and (P/S)_vitrified_ are measured values (1.12 and 2.2 respectively). GDH is a hexamer with a total molecular weight of 318 kD. There are two NADH binding sites per subunit, one with a K_D_ of ∼20 μM and a second with a K_D_ of ∼50 μM^5^. Our model is returning an average K_D_ value. In the absence of glutamate, GDH was shown to bind 7-8 molecules per hexamer^6^, and we assumed 8 binding sites per hexamer in the above kinetic analysis to estimate the concentration of binding sites from the protein concentration measured by UV spectroscopy (A280). In the model and data in Figure 2a and 2b in main test, [GDH]_T_ is the protein concentration x 8. The simulated line shown in Figure 2c in the main text was calculated as 1-f([GDH-NADH]) with [GDH] constant at 100 μM and [NADH] varied from 20 to 300 μM assuming a K_D_ of 26 μM.

#### 2.3 Robustness and reproducibly of FPRM of cellular NADH

As a ratiometric method, FPRM is independent of excitation intensity in the absence of pixel saturation, PMT gain (when changed by equal ratios for the two PMTs) and differences between objective lenses. Polarization ratio differences between objective lenses (Figure SI-2.3.1a) are due to intrinsic birefringence differences and increasing Z-axis polarization at the focus with increasing numerical aperture. The extent of z-polarization at the focus is shown in Figures SI-2.3.1b which shows the extent of z-polarized light along the Z axis for a 1.4 NA objective, and in Si-2.3.1c where the fractional X, Y and Z-polarized intensities are plotted as a function of numerical aperture (assuming X polarization at the back aperture of the objective). The normalization (Eq. SI-2.2) used to estimate the fractional unbound NADH allows for direct comparison of results between experiments and instruments as long as the normalization values (S/P_free_ and S/P_vitrified_) are measured for each system. In general, it is best to measure these two factors before the start of each experiment.

**Figure SI-2.3.1.**
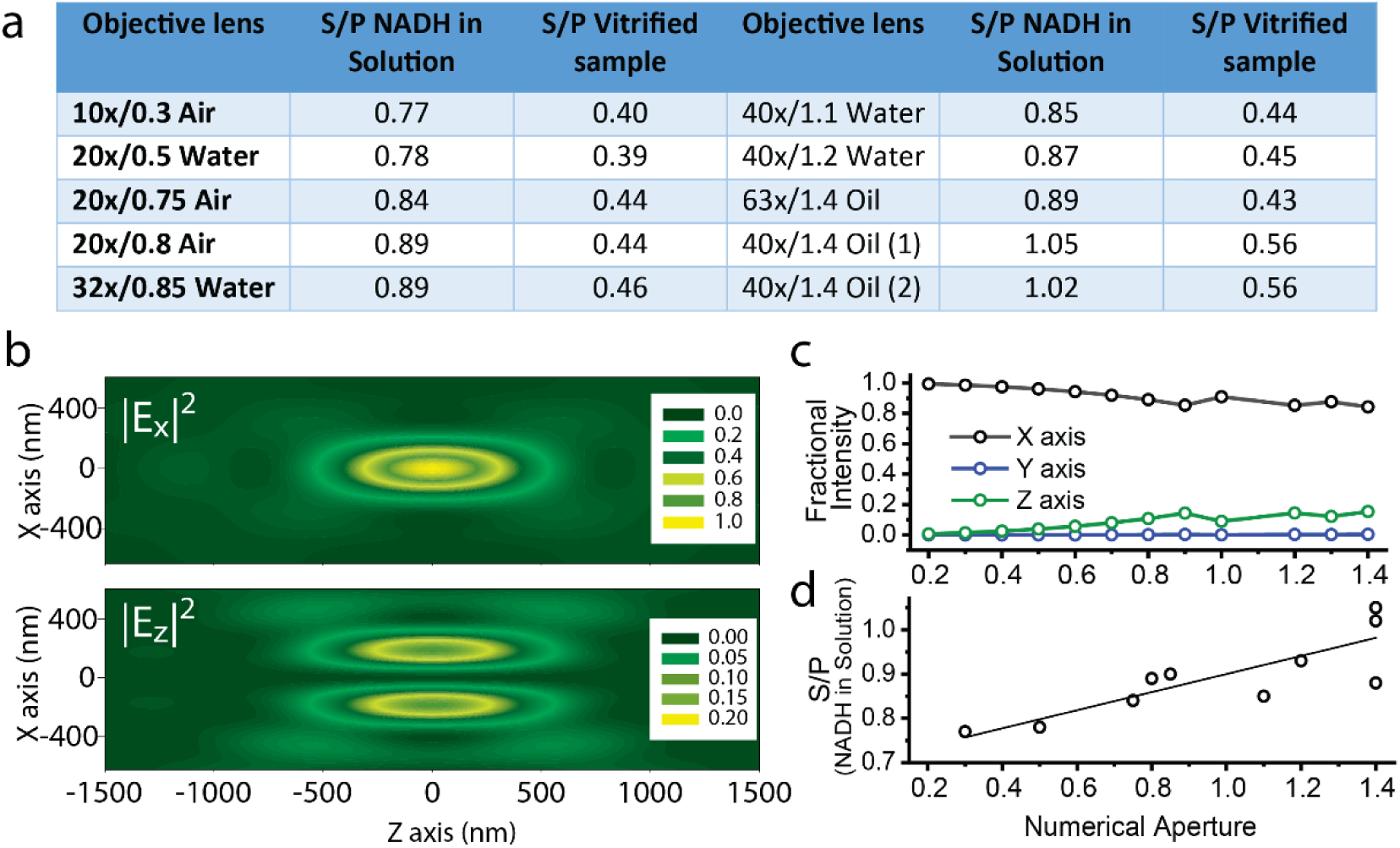
FPRM optical considerations. **(a)** Different objective lenses produce differing S/P ratios due to NA-dependent z-polarization component at focus and design-dependent birefringence. **(b)** Point spread functions of the X-polarized excitation (top) and the Z-polarized excitation for a 1.4 NA objective. **(c)** Fractional intensities of the X, Y and Z components at the focus as a function numerical aperture. Calculations were carried using the integral vector approach of Richards and Wolf^7^. (Note – the two “bumps” in the calculated polarization traces Figure SI-2.3.1c at NA=1.0 and NA = 1.3 result from changing the index of refraction from air, to water to oil.). **(d)** In general, the S/P ratio increases with numerical aperture as expected due to loss of X-polarized excitation as the NA increases.

FPRM is independent of laser power ss long as the S and P pixel values are not saturated. Note that saturation can be obfuscated by averaging. If averaging, the unaveraged pixels values must also be unsaturated.

**Figure SI-2.3.1.**
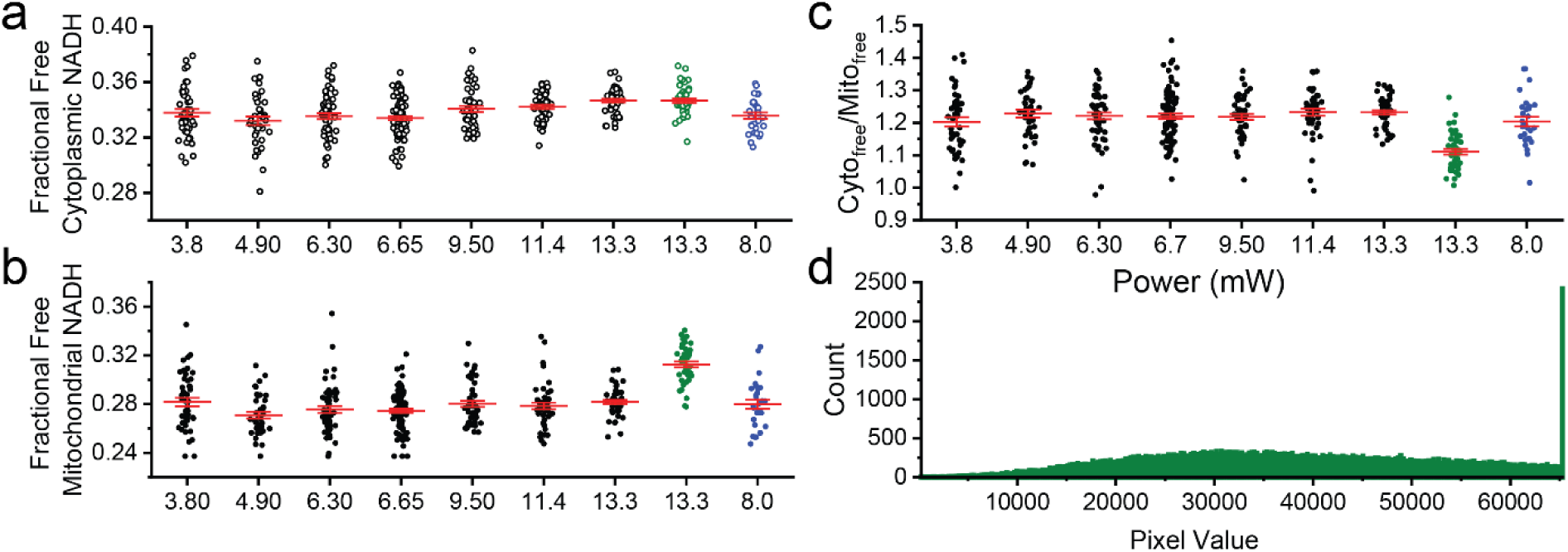
**(a)-(c)** FPRM parameters from MDA-MB-231 cells measured as a function of 730 nm laser power at the sample delivered through a 63x/1.4 NA objective lens. Different fields of view of the same dish of cells were used for each image analyzed. PMT gains (HV) were set to 900V for focal plane powers 3.8 – 6.7 mW and 800V for 9.5-13.3 mW to avoid saturating pixels at the higher power. Saturation will cause artifactual higher S/P ratios for the mitochondrial compartment since the P_mito_ region pixels are the brightest and will be the first to saturate, as seen in the 13.3 mW @ 900v PMT HV mitochondrial values which are the green 13.3 mW data points. Blue data points are FPRM parameters taken using a 40x/1.4 objective at 8 mW. The 40x oil objective has a significantly different free and vitrified S/P normalization values from the 63x oil objective, yet when normalized that cell data yields similar FPRM parameter values to those measured with the 63x objective. **(d)** Pixel value histogram of the mitochondrial regions from the green data points. Pixel saturation causes erroneously smaller S/P values from the mitochondrial regions, resulting in an overestimation of the free NADH. Images: 512×512, 1.54 μs pixel time. All of the FPRM images are 16-bit data, line averaged 4 times.

In addition to acquisition and optical factors discussed above, image segmentation parameters used for compartment separation could also result in different values for NADH_free_ in the two compartments and the Cyto/Mito ratio. We use mitochondrial segmentation by thresholding and find that there is a broad range of threshold values that still return statistically similar results, as shown in Figure SI-2.3.2a and b. As expected, there is a decrease in the estimate of cytoplasmic free NADH as segmentation threshold values increase and more of the mitochondrial compartment pixels are included as part of the cytoplasm mask.

**Figure SI-2.3.2.**
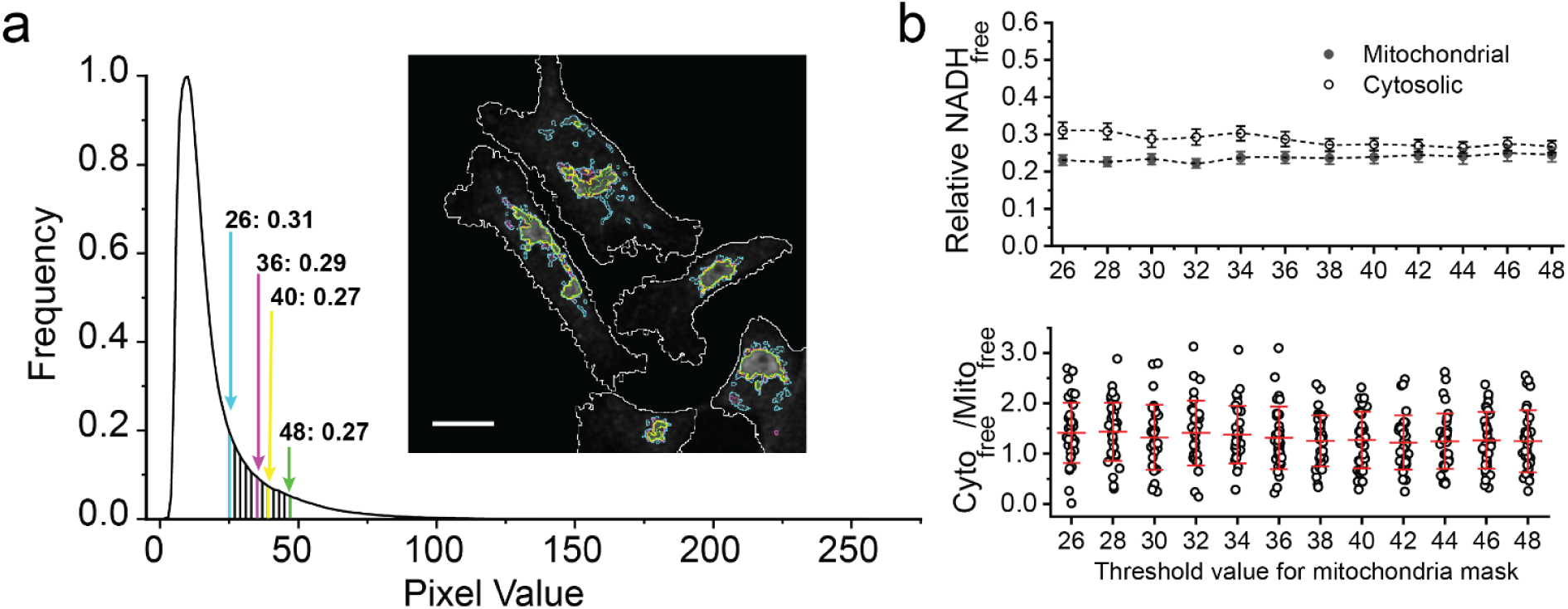
Sensitivity of FPRM to differing thresholding values. An FPRM image of MDA-MB-231 cells was analyzed using different mitochondrial pixel thresholds. **(a)** Histogram of the pixel intensities from within the cell boundaries. Color-coded segmentation regions are shown in the image insert (cropped from the full image of 37 cells) and indicated on the histogram. The image used for mask creation was the summed S and P intensity image (32-bit float) converted to 8-bit before thresholding to create the segmentation masks. NADH_free_ values are calculated from the original 16-bit images. **(b)** FPRM parameters from the full field of view as a function of threshold values from 26 to 48. (Upper plot – error bars are SEM, lower scatter plot – red bars are SD.) Although all of the values are statistically significant based on P-values (due to the large standard deviations from cell-cell heterogeneity), the free cytoplasmic NADH value does decrease at the highest threshold values as more of the mitochondria edges are included in the S/P value of the cytoplasmic compartment.

### Section SI-3. Stress from prolonged two-photon imaging in the 700 nm range

**Figure SI-3.**
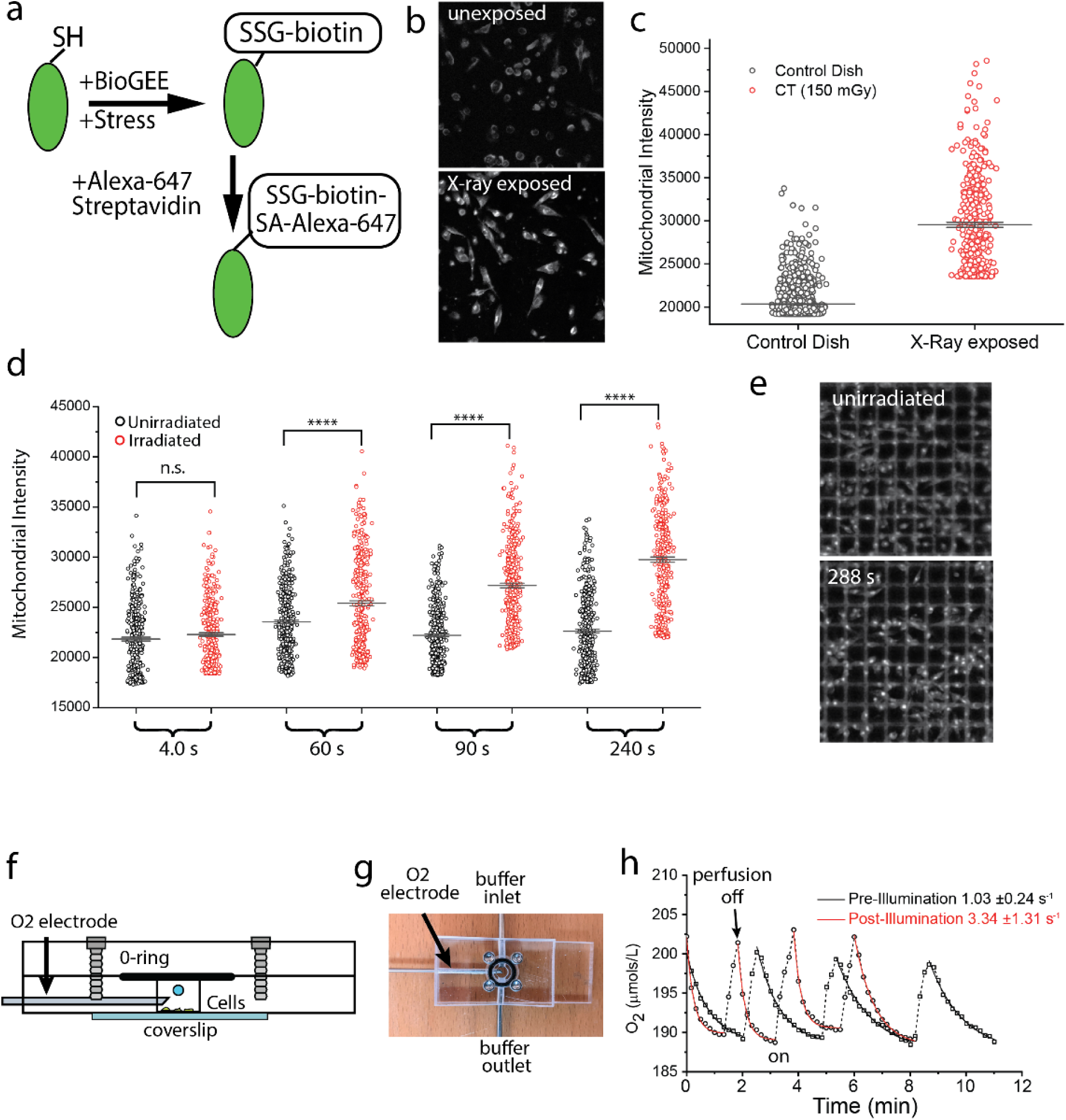
Femtosecond pulsed excitation in the 700 nm range causes oxidative stress. **(a)** Biotinylated glutathione ethyl ester (BioGEE) is a cell-permeant, biotinylated glutathione analog used to detect glutathiolation of proteins during oxidative stress. MDA-MB-231 cells were preincubated for 1 hour with BioGEE (250 μM) in Hepes buffered media, and then exposed to oxidative stress. After exposure, cells are permeabilized (4% PEG-8000, 1mM EGTA, 10 mM PIPES pH 7.2 and 0.2% Triton) and incubated with fluorescent streptavidin. The extent of protein glutathiolation can be quantified using fluorescence microscopy. **(b, c):** Verification of the assay using stress caused by X-ray exposure. BioGEE-loaded cells in a 35 mm cell culture dish were placed in a micro-CT (Bruker Skyscan 1276) and exposed to 150 mGr X-ray dose, after which the exposed dish and a +BioGee control dish were stained with Alexa-647 streptavidin and imaged by confocal microscopy (b). **(c)** Mean pixel values from mitochondrial regions indicate increased streptavidin staining after X-ray exposure compared to the unexposed cells. Increases in streptavidin signal is found primarily in the mitochondria. (d, e) Mean mitochondrial pixel values after repeated scans using at 11 mW of 700 nm femtosecond (∼120 fs) pulsed irradiation delivered through a Zeiss C-Achroplan 32x/0.85 W objective. The pixel dwell time was 2 μs. Cells were plated on gridded dishes, loaded with BioGee and a 2×3 tile scan was carried out (512×512 frames, 1.54 μs pixel time, 1.0 s per frame) for 4, 60, 90 and 240 frames. Control values in **(d)** are from regions of the gridded dishes not exposed to 700 nm excitation. For FPRM imaging, we typically use an average each line 4 times. **(f, g)** Respiration rates are also affected by prolonged scanning. MCF10a cells were plated on a coverslip glued to a lab-built Plexiglas chamber which has ports for fresh buffer and a phosphorescence-based oxygen probe (NeoFox Oxygen probe, Ocean Optics, Orlando, FL, USA). **(h)** Ambient oxygenated imaging buffer (∼210 μmol/L O2) is perfused through the imaging chamber and when perfusion is stopped a decrease in the O2 is seen due to respiration. The rate of change reflects the respiration rate of the cells. The relative respiration rate increased 3-fold after 200 seconds of scanning the cells with 10 mW of at 730 nm pulsed irradiation delivered through a 20x/1.0 NA objective lens.

### Section SI-4. Elaboration of Raman Data

**SI-4.**
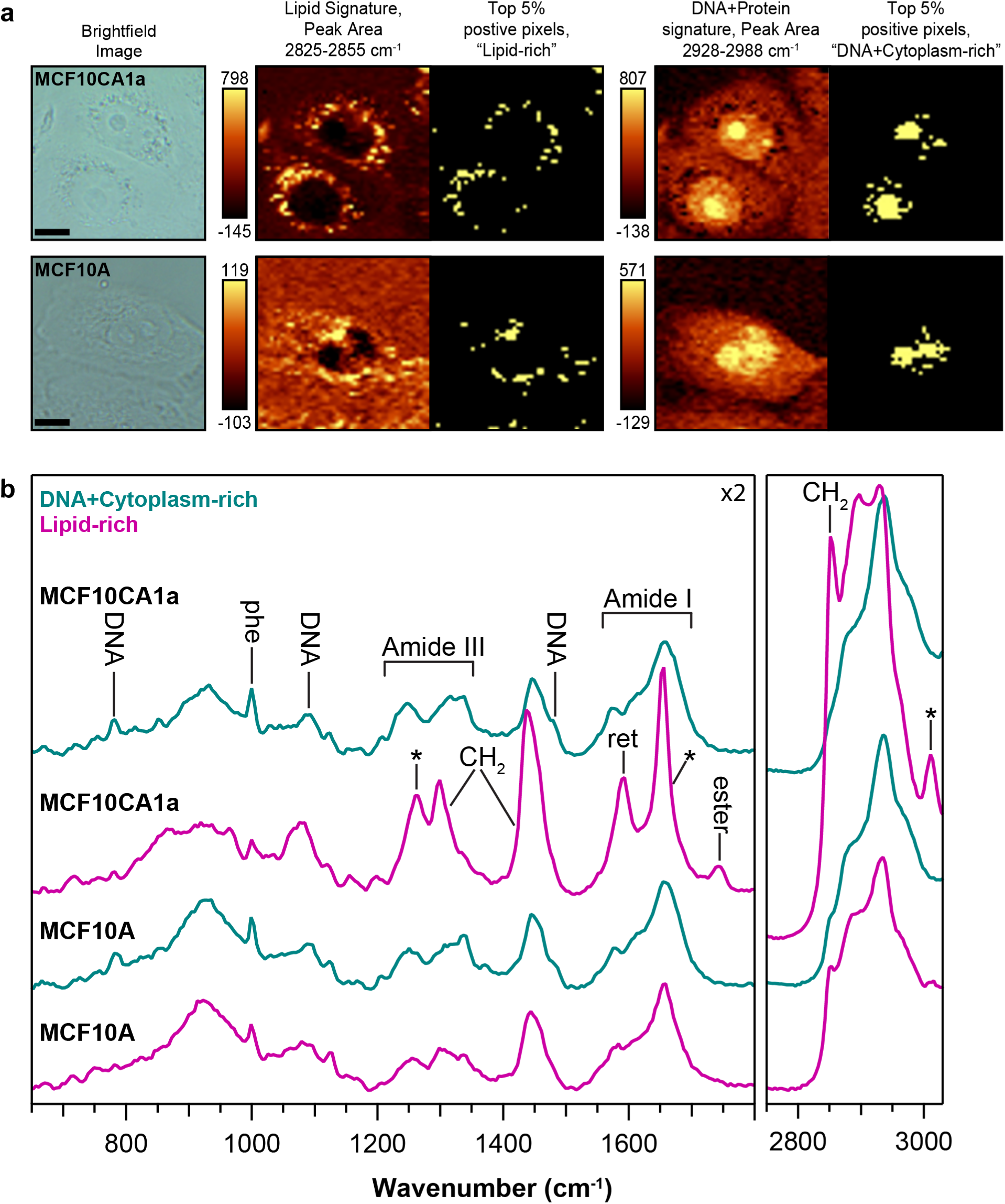
Raman mapping of MCF10CA1a and MCF10A. (a) Brightfield images are shown on the far left corresponding to the Raman mapping regions for the two cell lines: MCF10CA1a (Top) and MCF10A (Bottom). Unadjusted peak area false color images (with spectral integration and area value ranges indicated) are shown paired with the top 5% positive-value pixels (yellow pixels) that were averaged to generate the component-rich Raman spectra in Fig 3d. (b) An enlarged view of the average Raman spectra from Fig. 3d with labeled peaks showing the fingerprint region (below 1800 cm^-1^, shown left at 2-times y-scale) and the CH-stretching peak envelope (shown right, 2900 cm^-1^ region). The DNA/cytoplasm-rich spectrum is consistent overall with general proteins including contributions from amide bonds (the peak envelopes of Amide I and III are roughly indicated) and amino acids (e.g., phenylalanine, labeled ‘phe’), among others. Characteristic DNA (and RNA) backbone and base-associated peaks are also evident (including 783 cm^-1^, 1090 cm^-1^, and 1491 cm^-1^)^8^. The lipid-rich averages, particularly that from CA1a, contain peaks characteristic of lipids (including 2850 cm^-1^, 1300 cm^-1^, among others), consistent with acylglycerols (with the ester bond peak at 1742 cm^-1^ and lacking the characteristic cholesterol peak at ∼700 cm^-1^). Protein contributions are also evident (spectra were only averaged and no form of spectral unmixing was performed). The MCF10A lipid-rich spectrum contains a lower lipid-relative-to-protein presence compared to CA1a and, qualitatively at the raw spectrum level, lipid signatures generally appeared less pronounced. The CA1a lipid peaks labeled with an asterisk (including ∼1265 cm^-1^, ∼1654 cm^-1^ and 3011 cm^-1^) are consistent with the presence of unsaturation in the probed lipid population36. Notably, the peak at 1594 cm^-1^ assigned as retinol-like, labeled ‘ret’ (for similar spectrum see retinyl palmitate in ref [9] ^9^ is absent in MCF10A. Scale Bar 10 µm.

### Section SI-5. Heterogeneity in MDA-MB-231 response to KCN, FCCP, 2DG and AZD7574 as detected by FPRM

**Figure SI-5a.**
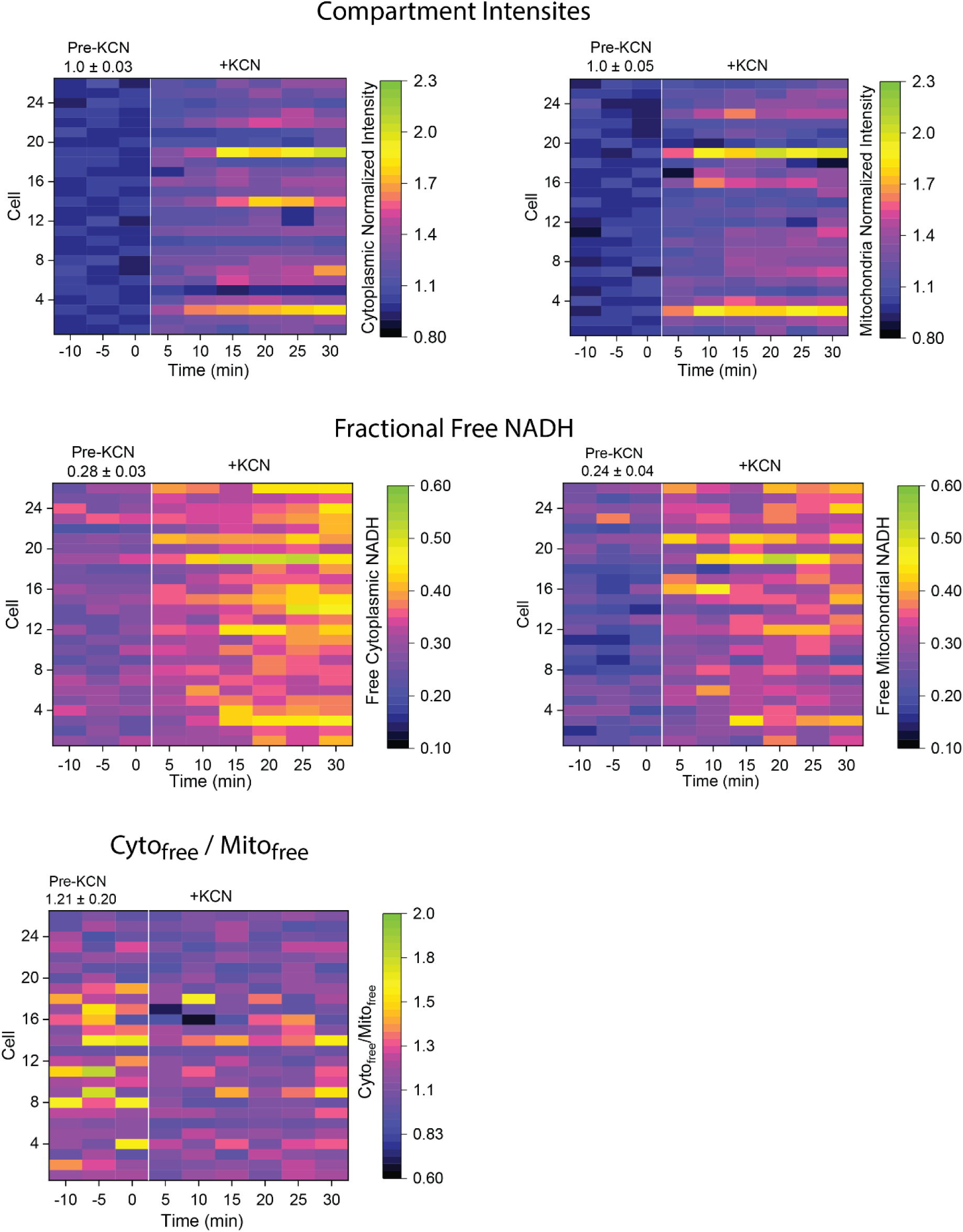
Single cell responses to KCN inhibition. FPRM parameters vs. time from 26 randomly selected cells from the KCN treatment data set shown in Figure 4. Individual cells were cut from the time series stack in ImageJ and analyzed.

**Figure SI-5b.**
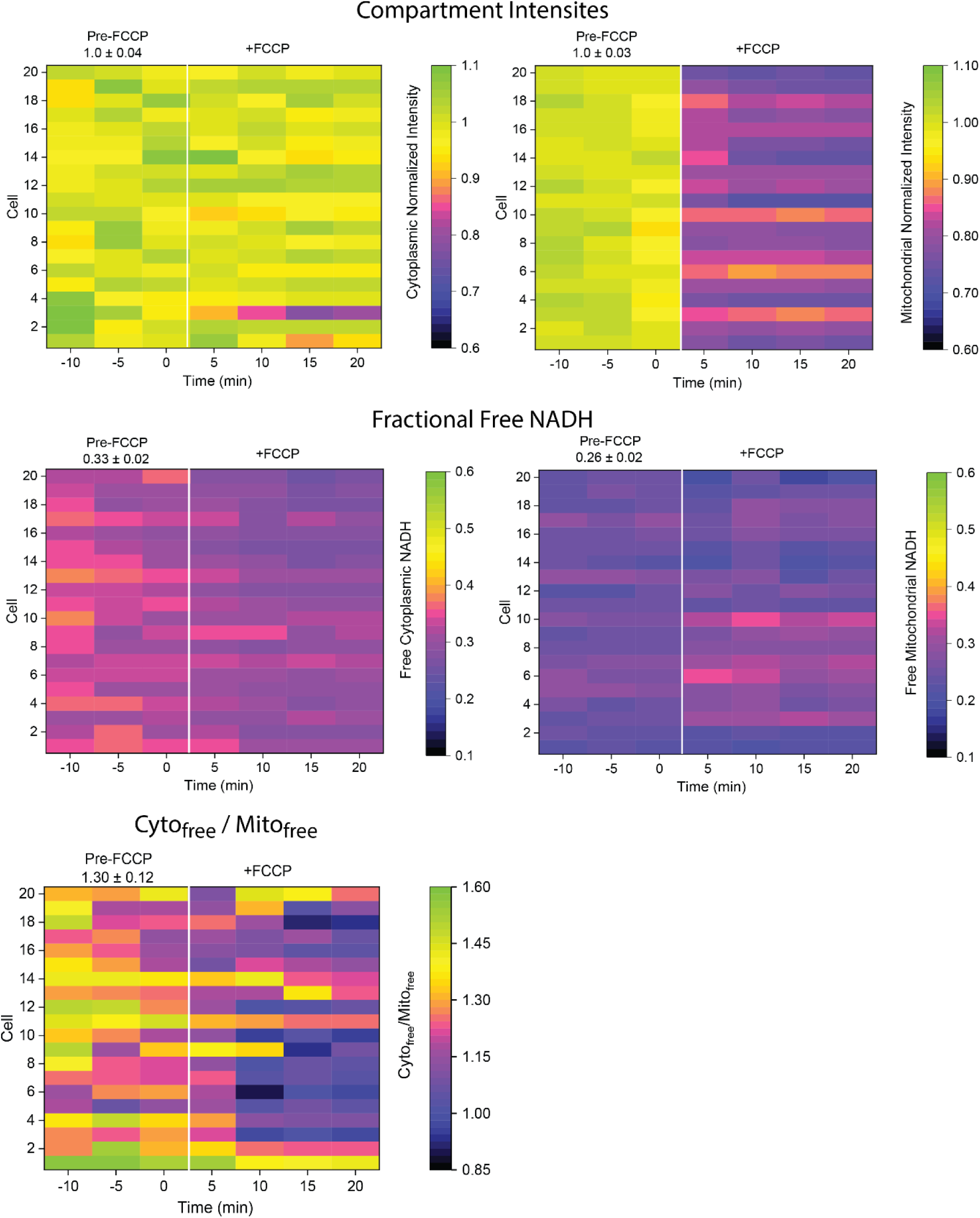
Single cell responses to FCCP inhibition. FPRM parameters vs. time from 20 randomly selected cells from the FCCP treatment data set shown in Figure 4. Individual cells were cut from the time series stack in ImageJ and analyzed.

**Figure SI-5c.**
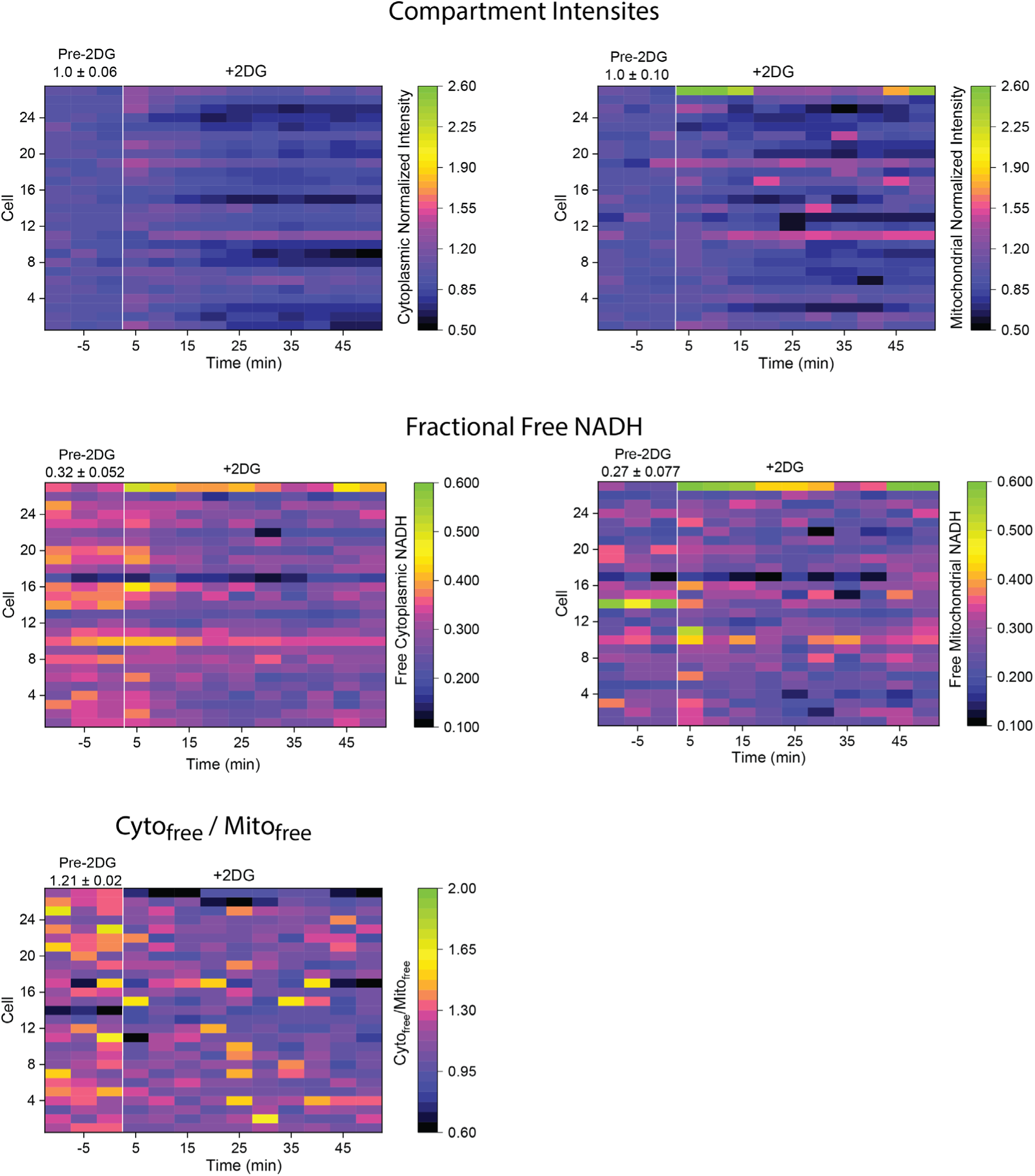
Single cell responses to 2DG inhibition. FPRM parameters vs. time from 27 randomly selected cells from the 2DG treatment data set shown in Figure 4. Individual cells were cut from the time series stack in ImageJ and analyzed.

**Figure SI-5d.**
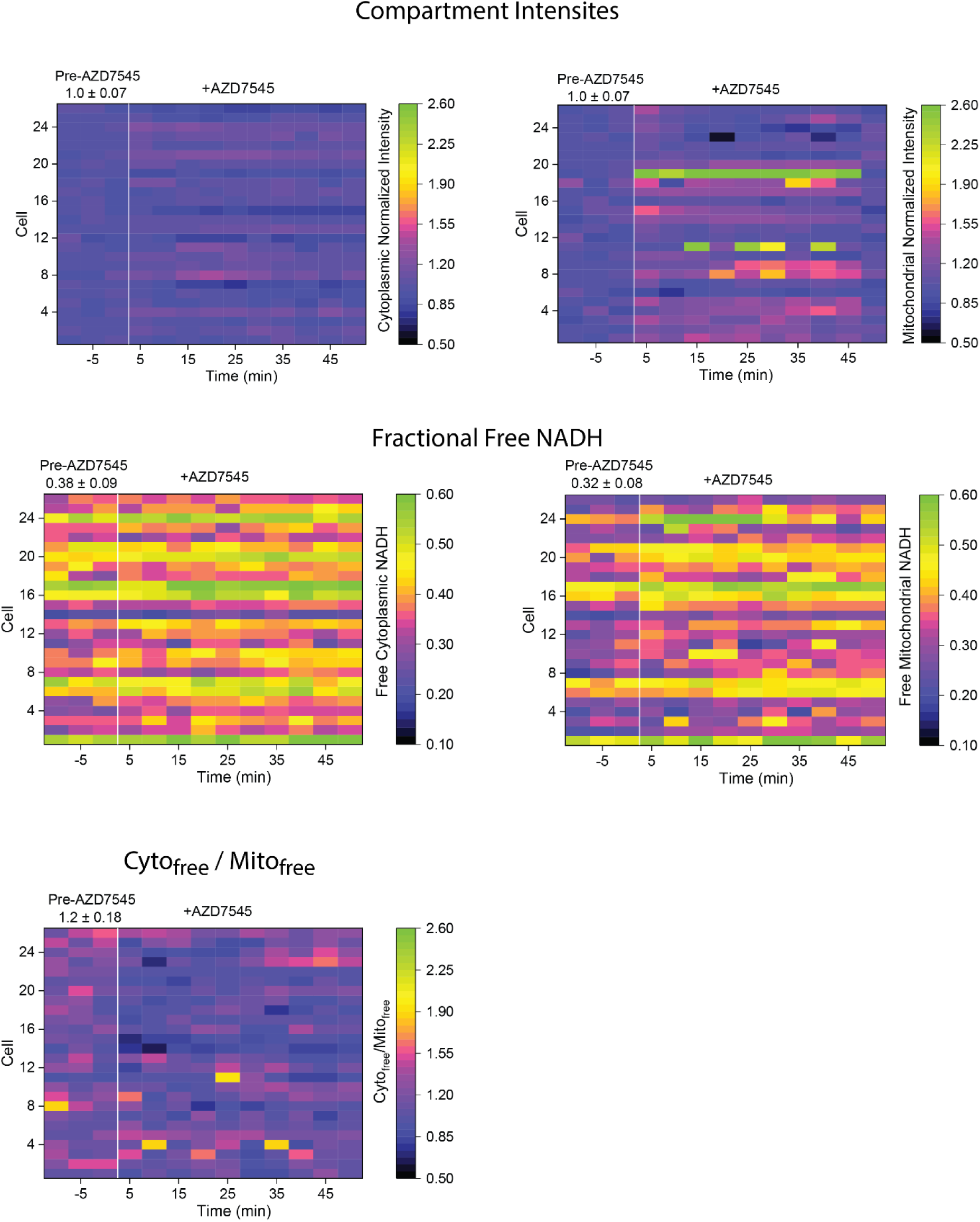
Single cell responses to AZD7475 inhibition. FPRM parameters vs. time from 26 randomly selected cells from the AZD7574 treatment data set shown in Figure 4. Individual cells were cut from the time series stack in ImageJ and analyzed.

### Section SI-6. Videos of FPRM changes during cell motility

**FastCell_Video.mp3: C**olor-coded compartmental NADH_free_ values and FPRM parameter plots from a rapidly migrating MDA-MD-323 cell.

**SlowCell_Video.mp3: C**olor-coded compartmental NADH_free_ values and FPRM parameter plots from a ∼stationary MDA-MD-323 cell.

**Appendix SI-Table 1.** Estimates of NADH and NADH binding enzyme concentrations in cells.

|  | Cytoplasm | Mitochondria |
| --- | --- | --- |
| $\text{NAD}^+/\text{NADH}$ Ratio | 60-1000<sup>10</sup> | 7-10<sup>10</sup> |
| Total $[\text{NAD}^+] + [\text{NADH}]$ | $\sim 350 \mu\text{M}^{10,11}$ | $\sim 500 \mu\text{M}^{10}$ |
| Free $[\text{NAD}^+]$ and $[\text{NADH}]$ | $\sim 50$ to $100 \mu\text{M}^{11}$ | 0 to $\sim 30 \mu\text{M}^{12}$ |
| Number of proteins with NADH binding sites (both compartments) | $\sim 60^{13,*}$ | |
| Range of $K_m$ ($\sim K_D$) values | $0.1\text{-}20 \mu\text{M}^{14,**}$ | |
| Concentration range of NADH binding proteins | $0.1\text{-}10 \mu\text{M}^{14}$ | |

\*Summarized from the BRENDA database with redundancies removed.
\*\* The majority the NADH binding enzymes in the BRENDA database where in this range.

**Appendix SI-Table 2.**
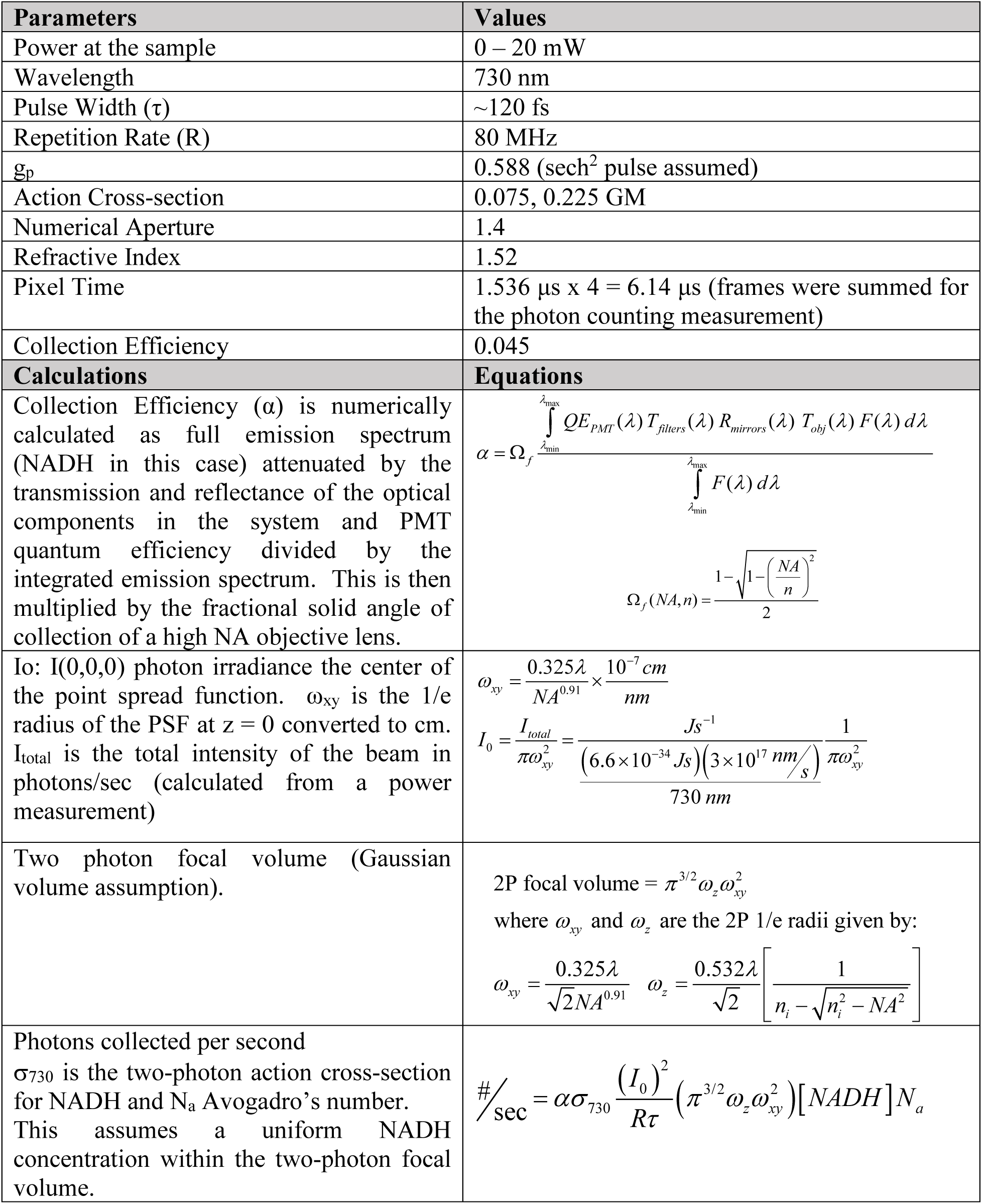
Information on the NADH imaging calculations shown in Figure SI-1b.

## References

1 Pavlova, N. N. & Thompson, C. B. The Emerging Hallmarks of Cancer Metabolism. Cell Metabolism 23, 27–47, doi:10.1016/j.cmet.2015.12.006 (2016).

2 Bergers, G. & Fendt, S. M. The metabolism of cancer cells during metastasis. Nature Reviews Cancer 21, 162–180, doi:10.1038/s41568-020-00320-2 (2021).

3 DeBerardinis, R. J. & Keshari, K. R. Metabolic analysis as a driver for discovery, diagnosis, and therapy. Cell 185, 2678–2689, doi:10.1016/j.cell.2022.06.029 (2022).

4 Hong, S., Pawel, G. T., Pei, R. & Lu, Y. Recent progress in developing fluorescent probes for imaging cell metabolites. Biomed Mater 16, doi:10.1088/1748-605X/abfd11 (2021).

5 Gooz, M. & Maldonado, E. N. Fluorescence microscopy imaging of mitochondrial metabolism in cancer cells. Front Oncol 13, 1152553, doi:10.3389/fonc.2023.1152553 (2023).

6 Chance, B. & Thorell, B. Fluorescence measurements of mitochondrial pyridine nucleotide in aerobiosis and anaerobiosis. Nature 184, 931–934, doi:10.1038/184931a0 (1959).

7 Shuttleworth, C. W. Use of NAD(P)H and flavoprotein autofluorescence transients to probe neuron and astrocyte responses to synaptic activation. Neurochemistry International 56, 379–386, 10.1016/j.neuint.2009.12.015 (2010).

8 Reiss, P. D., Zuurendonk, P. F. & Veech, R. L. Measurement of tissue purine, pyrimidine, and other nucleotides by radial compression high-performance liquid chromatography. Analytical Biochemistry 140, 162–171, 10.1016/0003-2697(84)90148-9 (1984).

9 Kasischke, K. A., Vishwasrao, H. D., Fisher, P. J., Zipfel, W. R. & Webb, W. W. Neural activity triggers neuronal oxidative metabolism followed by astrocytic glycolysis. Science 305, 99–103, doi:10.1126/science.1096485 (2004).

10 Piston, D. W., Masters, B. R. & Webb, W. W. Three-dimensionally resolved NAD(P)H cellular metabolic redox imaging of the in situ cornea with two-photon excitation laser scanning microscopy. J Microsc 178, 20–27, doi:10.1111/j.1365-2818.1995.tb03576.x (1995).

11 Bird, D. K. et al. Metabolic mapping of MCF10A human breast cells via multiphoton fluorescence lifetime imaging of the coenzyme NADH. Cancer Res 65, 8766–8773, doi:10.1158/0008-5472.CAN-04-3922 (2005).

12 Blacker, T. S. et al. Separating NADH and NADPH fluorescence in live cells and tissues using FLIM. Nat Commun 5, 3936, doi:10.1038/ncomms4936 (2014).

13 Evers, M. et al. Enhanced quantification of metabolic activity for individual adipocytes by label-free FLIM. Sci Rep 8, 8757, doi:10.1038/s41598-018-27093-x (2018).

14 Liu, Z. et al. Mapping metabolic changes by noninvasive, multiparametric, high-resolution imaging using endogenous contrast. Science Advances 4, doi:10.1126/sciadv.aap9302 (2018).

15 Skala, M. C. et al. In vivo multiphoton microscopy of NADH and FAD redox states, fluorescence lifetimes, and cellular morphology in precancerous epithelia. Proceedings of the National Academy of Sciences of the United States of America 104, 19494–19499, doi:10.1073/pnas.0708425104 (2007).

16 Vergen, J. et al. in Microscopy and Microanalysis Vol. 18 761–770 (2012).

17 Walsh, A. J. et al. Classification of T-cell activation via autofluorescence lifetime imaging. Nat Biomed Eng 5, 77–88, doi:10.1038/s41551-020-0592-z (2021).

18 Poudel, C., Mela, I. & Kaminski, C. F. High-throughput, multi-parametric, and correlative fluorescence lifetime imaging. Methods and Applications in Fluorescence 8, 024005, doi:10.1088/2050-6120/ab7364 (2020).

19 Xu, C. & Zipfel, W. R. Multiphoton excitation of fluorescent probes. Cold Spring Harbor protocols 2015 3, 250–258 (2015).

20 Vishwasrao, H. D., Heikal, A. A., Kasischke, K. A. & Webb, W. W. Conformational dependence of intracellular NADH on metabolic state revealed by associated fluorescence anisotropy. J Biol Chem 280, 25119–25126, doi:10.1074/jbc.M502475200 (2005).

21 Smith, H. E. et al. The use of NADH anisotropy to investigate mitochondrial cristae alignment. Sci Rep 14, 5980, doi:10.1038/s41598-024-55780-5 (2024).

22 Yu, Q. & Heikal, A. A. Two-photon autofluorescence dynamics imaging reveals sensitivity of intracellular NADH concentration and conformation to cell physiology at the single-cell level. Journal of Photochemistry and Photobiology B: Biology 95, 46–57, doi:10.1016/j.jphotobiol.2008.12.010 (2009).

23 Zheng, W., Li, D. & Qu, J. Y. Monitoring changes of cellular metabolism and microviscosity in vitro based on time-resolved endogenous fluorescence and its anisotropy decay dynamics. J Biomed Opt 15, 037013, doi:10.1117/1.3449577 (2010).

24 Churchich, J. E. The binding of NADH by glutamate dehydrogenase as measured by polarization of fluorescence. Biochim Biophys Acta 147, 32–38, doi:10.1016/0005-2795(67)90086-4 (1967).

25 Malcolm, A. D. B. Coenzyme Binding to Glutamate Dehydrogenase. European Journal of Biochemistry 27, 453–461, 10.1111/j.1432-1033.1972.tb01860.x (1972).

26 Lakowicz, J. R. Principles of fluorescence spectroscopy. 2nd edn, (Kluwer Academic/Plenum, 1999).

27 Xie, N. et al. NAD+ metabolism: pathophysiologic mechanisms and therapeutic potential. Signal Transduction and Targeted Therapy 5, doi:10.1038/s41392-020-00311-7 (2020).

28 Tan, M. L. et al. Endothelial cells metabolically regulate breast cancer invasion toward a microvessel. APL Bioeng 7, 046116, doi:10.1063/5.0171109 (2023).

29 Cambronne, X. A. et al. Biosensor reveals multiple sources for mitochondrial NAD+. Science 352, 1474–1477, doi:10.1126/science.aad5168 (2016).

30 Carles Cantó, K. M., Johan Auwerx. NAD+ metabolism and the control of energy homeostasis - a balancing act between mitochondria and the nucleus. Cell Metabolism 22, 31–53, doi:10.1016/j.cmet.2015.05.023.NAD (2015).

31 Hu, Q. et al. Genetically encoded biosensors for evaluating NAD+/NADH ratio in cytosolic and mitochondrial compartments. Cell Reports Methods 1, 100116, doi:10.1016/j.crmeth.2021.100116 (2021).

32 Sullivan, D. M., Levine, R. L. & Finkel, T. Detection and affinity purification of oxidant-sensitive proteins using biotinylated glutathione ethyl ester. Methods in Enzymology 353, 101–113, doi:10.1016/S0076-6879(02)53040-8 (2002).

33 Santner, S. J. et al. in Breast Cancer Research and Treatment. 101–110.

34 Vidavsky, N. et al. Mapping and Profiling Lipid Distribution in a 3D Model of Breast Cancer Progression. ACS Cent Sci 5, 768–780, doi:10.1021/acscentsci.8b00932 (2019).

35 Kunitake, J. A. M. R. et al. Biomineralogical signatures of breast microcalcifications. Science Advances 9, eade3152, doi:doi:10.1126/sciadv.ade3152 (2023).

36 Czamara, K. et al. Raman spectroscopy of lipids: a review. Journal of Raman Spectroscopy 46, 4–20, 10.1002/jrs.4607 (2015).

37 Movasaghi, Z., Rehman, S. & Rehman, I. U. Raman Spectroscopy of Biological Tissues. Applied Spectroscopy Reviews 42, 493–541, doi:10.1080/05704920701551530 (2007).

38 Zipfel, W. R. et al. Live tissue intrinsic emission microscopy using multiphoton-excited native fluorescence and second harmonic generation. Proceedings of the National Academy of Sciences 100, 7075–7080, doi:doi:10.1073/pnas.0832308100 (2003).

39 Benz, R. & McLaughlin, S. The molecular mechanism of action of the proton ionophore FCCP (carbonylcyanide p-trifluoromethoxyphenylhydrazone). Biophys J 41, 381–398, doi:10.1016/s0006-3495(83)84449-x (1983).

40 Hu, Q. et al. Genetically encoded biosensors for evaluating NAD(+)/NADH ratio in cytosolic and mitochondrial compartments. Cell Rep Methods 1, doi:10.1016/j.crmeth.2021.100116 (2021).

41 Zhang, D. et al. 2-Deoxy-D-glucose targeting of glucose metabolism in cancer cells as a potential therapy. Cancer Lett 355, 176–183, doi:10.1016/j.canlet.2014.09.003 (2014).

42 Cheng, G. et al. Mitochondria-targeted drugs synergize with 2-deoxyglucose to trigger breast cancer cell death. Cancer Res 72, 2634–2644, doi:10.1158/0008-5472.Can-11-3928 (2012).

43 Luengo, A. et al. Increased demand for NAD+ relative to ATP drives aerobic glycolysis. Molecular Cell 81, 691–707.e696, doi:10.1016/j.molcel.2020.12.012 (2021).

44 Wengrowski, A. M., Kuzmiak-Glancy, S., Jaimes, R., 3rd & Kay, M. W. NADH changes during hypoxia, ischemia, and increased work differ between isolated heart preparations. Am J Physiol Heart Circ Physiol 306, H529–537, doi:10.1152/ajpheart.00696.2013 (2014).

45 Shimpi, A. A. et al. Convergent Approaches to Delineate the Metabolic Regulation of Tumor Invasion by Hyaluronic Acid Biosynthesis. Adv Healthc Mater 12, e2202224, doi:10.1002/adhm.202202224 (2023).

46 Seo, B. R. et al. Collagen Microarchitecture Mechanically Controls Myofibroblast Differentiation. Proceedings of the National Academy of Sciences 117, doi:10.1073/pnas.1919394117 (2020).

47 Nguyen-Ngoc, K.-V. et al. ECM microenvironment regulates collective migration and local dissemination in normal and malignant mammary epithelium. Proceedings of the National Academy of Sciences 109, E2595–E2604, doi:10.1073/pnas.1212834109 (2012).

48 Winkler, J., Abisoye-Ogunniyan, A., Metcalf, K. J. & Werb, Z. Concepts of extracellular matrix remodelling in tumour progression and metastasis. Nature Communications 11, 1–19, doi:10.1038/s41467-020-18794-x (2020).

49 Paszek, M. J. et al. Tensional homeostasis and the malignant phenotype. Cancer Cell 8, 241–254, doi:10.1016/j.ccr.2005.08.010 (2005).

50 Sahai, E. & Marshall, C. J. in NATURE CELL BIOLOGY.

51 Ling, L. et al. Obesity-associated Adipose Stromal Cells Promote Breast Cancer Invasion Through Direct Cell Contact and ECM Remodeling. Adv Funct Mater 30, doi:10.1002/adfm.201910650 (2020).

52 Ishizaki, T. et al. Pharmacological properties of Y-27632, a specific inhibitor of rho-associated kinases. Mol Pharmacol 57, 976–983 (2000).

53 Guerra, F. S., Oliveira, R. G., Fraga, C. A. M., Mermelstein, C. D. S. & Fernandes, P. D. ROCK inhibition with Fasudil induces beta-catenin nuclear translocation and inhibits cell migration of MDA-MB 231 human breast cancer cells. Sci Rep 7, 13723, doi:10.1038/s41598-017-14216-z (2017).

## Supplementary Information References

1 Gorbunova, I. A. et al. Determination of fluorescence quantum yields and decay times of NADH and FAD in water–alcohol mixtures: The analysis of radiative and nonradiative relaxation pathways. Journal of Photochemistry and Photobiology A: Chemistry 436, 114388, 10.1016/j.jphotochem.2022.114388 (2023).

2 Xu, C. & Zipfel, W. R. Multiphoton excitation of fluorescent probes. Cold Spring Harbor protocols 2015 3, 250–258 (2015).

3 Köllner, M. & Wolfrum, J. How many photons are necessary for fluorescence-lifetime measurements? Chemical Physics Letters 200, 199–204, 10.1016/0009-2614(92)87068-Z (1992).

4 Smith, H. E. et al. The use of NADH anisotropy to investigate mitochondrial cristae alignment. Sci Rep 14, 5980, doi:10.1038/s41598-024-55780-5 (2024).

5 Delabar, J. M., Martin, S. R. & Bayley, P. M. The Binding of NADH and NADPH to Bovine-Liver Glutamate Dehydrogenase. European Journal of Biochemistry 127, 367–374, 10.1111/j.1432-1033.1982.tb06881.x (1982).

6 Shafer, J. A. et al. Binding of Reduced Cofactor to Glutamate Dehydrogenase. European Journal of Biochemistry 31, 166–171, 10.1111/j.1432-1033.1972.tb02515.x (1972).

7 Richards, B. & Wolf, E. Electromagnetic diffraction in optical systems, II. Structure of the image field in an aplanatic system. Proceedings of the Royal Society of London. Series A. Mathematical and Physical Sciences 253, 358–379, doi:doi:10.1098/rspa.1959.0200 (1959).

8 Peticolas, W. L. Applications of Raman spectroscopy to biological macromolecules. Biochimie 57, 417–428, 10.1016/S0300-9084(75)80328-2 (1975).

9 Abramczyk, H., Imiela, A. & Surmacki, J. Novel strategies of Raman imaging for monitoring intracellular retinoid metabolism in cancer cells. Journal of Molecular Liquids 334, 116033, 10.1016/j.molliq.2021.116033 (2021).

10 Stein, L. R. & Imai, S.-i. The dynamic regulation of NAD metabolism in mitochondria. Trends in Endocrinology & Metabolism 23, 420–428, 10.1016/j.tem.2012.06.005 (2012).

11 Cambronne, X. A. et al. Biosensor reveals multiple sources for mitochondrial NAD+. Science 352, 1474–1477, doi:10.1126/science.aad5168 (2016).

12 Zhao, Y. et al. Genetically encoded fluorescent sensors for intracellular NADH detection. Cell Metab 14, 555–566, doi:10.1016/j.cmet.2011.09.004 (2011).

13 Chang, A. et al. BRENDA, the ELIXIR core data resource in 2021: new developments and updates. Nucleic Acids Research 49, D498–D508, doi:10.1093/nar/gkaa1025 (2020).

14 Albe, K. R., Butler, M. H. & Wright, B. E. Cellular concentrations of enzymes and their substrates. J Theor Biol 143, 163–195, doi:10.1016/s0022-5193(05)80266-8 (1990).

